# Single Cell RNA-seq and Mass Cytometry Reveals a Novel and a Targetable Population of Macrophages in Idiopathic Pulmonary Fibrosis

**DOI:** 10.1101/2021.01.04.425268

**Authors:** EA Ayaub, S Poli, J Ng, T Adams, J Schupp, L Quesada-Arias, F Poli, C Cosme, M Robertson, J Martinez-Manzano, X Liang, J Villalba, J Lederer, SG Chu, BA Raby, G Washko, C Coarfa, MA Perrella, S El-Chemaly, N Kaminski, IO Rosas

## Abstract

In this study, we leveraged a combination of single cell RNAseq, cytometry by time of flight (CyTOF), and flow cytometry to study the biology of a unique macrophage population in pulmonary fibrosis. Using the profiling data from 312,928 cells derived from 32 idiopathic pulmonary fibrosis (IPF), 29 healthy control and 18 chronic obstructive pulmonary disease (COPD) lungs, we identified an expanded population of macrophages in IPF that have a unique transcriptional profile associated with pro-fibrotic signature. These macrophages attain a hybrid transitional state between alveolar and interstitial macrophages, are enriched with biological processes of pro-fibrotic immune cells, and express novel surface markers and genes that have not been previously reported. We then applied single cell CyTOF to simultaneously measure 37 markers to precisely phenotype the uniquely expanded macrophage subset in IPF lungs. The SPADE algorithm independently identified an expanded macrophage cluster, and validated CD84 and CD36 as novel surface markers that highly label this cluster. Using a separate validation cohort, we confirmed an increase in CD84^++^CD36^++^ macrophage population in IPF compared to control and COPD lungs by flow cytometry. Further, using the signature from the IPF-specific macrophages and the LINCS drug database, we predicted small molecules that could reverse the signature of IPF-specific macrophages, and validated two molecules, CRT and Cucur, using THP-1 derived human macrophages and precision-cut lung slices (PCLS) from IPF patients. Utilizing a multi-dimensional translational approach, our work identified a novel and targetable population of macrophages found in end-stage pulmonary fibrosis.

**One Sentence Summary:** Single cell RNAseq, CyTOF, and flow cytometry reveal the presence of an aberrant macrophage population in pulmonary fibrosis

## INTRODUCTION

Idiopathic pulmonary fibrosis (IPF) is a chronic, progressive, fibrosing interstitial pneumonia of unknown cause occurring primarily in older adults, with a median survival of 3 years after the initial diagnosis(1). Currently, lung transplantation is the only intervention that offers a mortality benefit in selected individuals. While there are two FDA approved therapies (2,3), these treatments do not significantly improve survival or quality of life. Therefore, there have been extensive efforts to better understand the pathobiology of IPF, the contributions of epithelial-mesenchymal cell interactions (4), and the role of immune cells in both initial and late stages of disease (5–7).

Macrophages are the most abundant immune cell type in the lungs(8), and have been increasingly recognized as major players in the initiation and development of the fibrotic response. These cells are highly plastic and can adapt to external stimuli by altering their functional phenotypes with multiple biological implications (9,10). These changes can be classified along the spectrum of the canonical M1 and M2 phenotypes(11), with characteristic secreted factors, surface markers and biological functions. Lung macrophages can additionally be classified based on their ontogeny (arising from the bone marrow, the embryonic yolk sac, or the fetal liver (12,13)), and their location within the lung (defining either Alveolar [AM] or Interstitial Macrophages [IM](14,15)). Recent scRNAseq studies have demonstrated the existence of a unique population of alveolar macrophages in pulmonary fibrosis lung explants(16–18), bronchoalveolar lavage (BAL) (17), and murine models of fibrotic lung disease(19,20), confirming macrophages as key players in the fibrotic response. The goal of this work is to phenotype the pro-fibrotic macrophages, their surface markers, and their interactions with other cells in in the fibrotic milieu and identify potential compounds that can modulate their behavior.

## RESULTS

### Myeloid cell subset and composition analysis of the different cell types

Our previously published work(18) on human lung explants from 32 IPF,18 COPD and 28 healthy controls was used for downstream analyses. From the complete scRNASeq dataset, myeloid cells were identified based on the gene expression of canonical markers for immune cells. A total of 216,978 cells were identified, comprising 109,820 interstitial macrophages (ITGAM^hi^), 60,478 alveolar macrophages (FABP4^hi^, C1QB^+^), 12,345 dendritic cells, 20,025 classical (CD14^+^) monocytes, and 14,110 non-classical (CD14^-^, CD16^+^) monocytes. Representative gene markers for each cell population described above are shown in a heatmap plot (Supplementary Figure 1A), along with Uniform Manifold Approximation and Projection (UMAP) plots of the myeloid compartment by cell population, disease state and individual subjects (Supplementary Figure 1B). Composition analyses for these myeloid cell subpopulations (Supplementary Figure 1C) revealed: 1) expansion of Interstitial Macrophages (IMΦ) in IPF lungs compared to COPD and Controls (*p-value <0.01*), 2) increased representation of Alveolar Macrophages (AMΦ) in both IPF and COPD when compared to Controls, but not statistically significant between the two disease conditions, and 3) increased abundance in both classical and non-classical monocytes in COPD lung explants than IPF and Controls(21) (*p-value <0.01*) (Supplementary Figure 1B-C).

### IPF-expanded macrophages (IPFeMΦ) are a discrete sub-population of cells with a pro-fibrotic and specific immunological signature

To achieve better clustering resolution of our subpopulations of interest, we limited the dataset further by excluding dendritic cells and Mast Cells and focusing on monocytes and macrophages. Low dimensional embeddings were made with the Potential of Heat-diffusion for Affinity-based Trajectory Embedding (PHATE) algorithm (22). The *slingshot* R package (23) was used to identify developmental trajectories and estimated pseudo-time distances. Regulons that were overexpressed across the pseudotime distance were reconstructed using the *pySCENIC* package (24). The PHATE embeddings (Figure 1A) revealed two different trajectories (T1 and T2), both of which begin with classical monocytes (T0). In the T1 trajectory, IPF-expanded macrophages (IPFeM**Φ**) comprised the distal branching structure, whereas FABP4+ Macrophages (FABP4+ MΦ) occupied the T2 trajectory. These findings provide evidence that the IPFeM**Φ** are a discrete cluster of cells with a transcriptional gene signature that differs from the other myeloid cell subpopulations.

**Figure 1:**
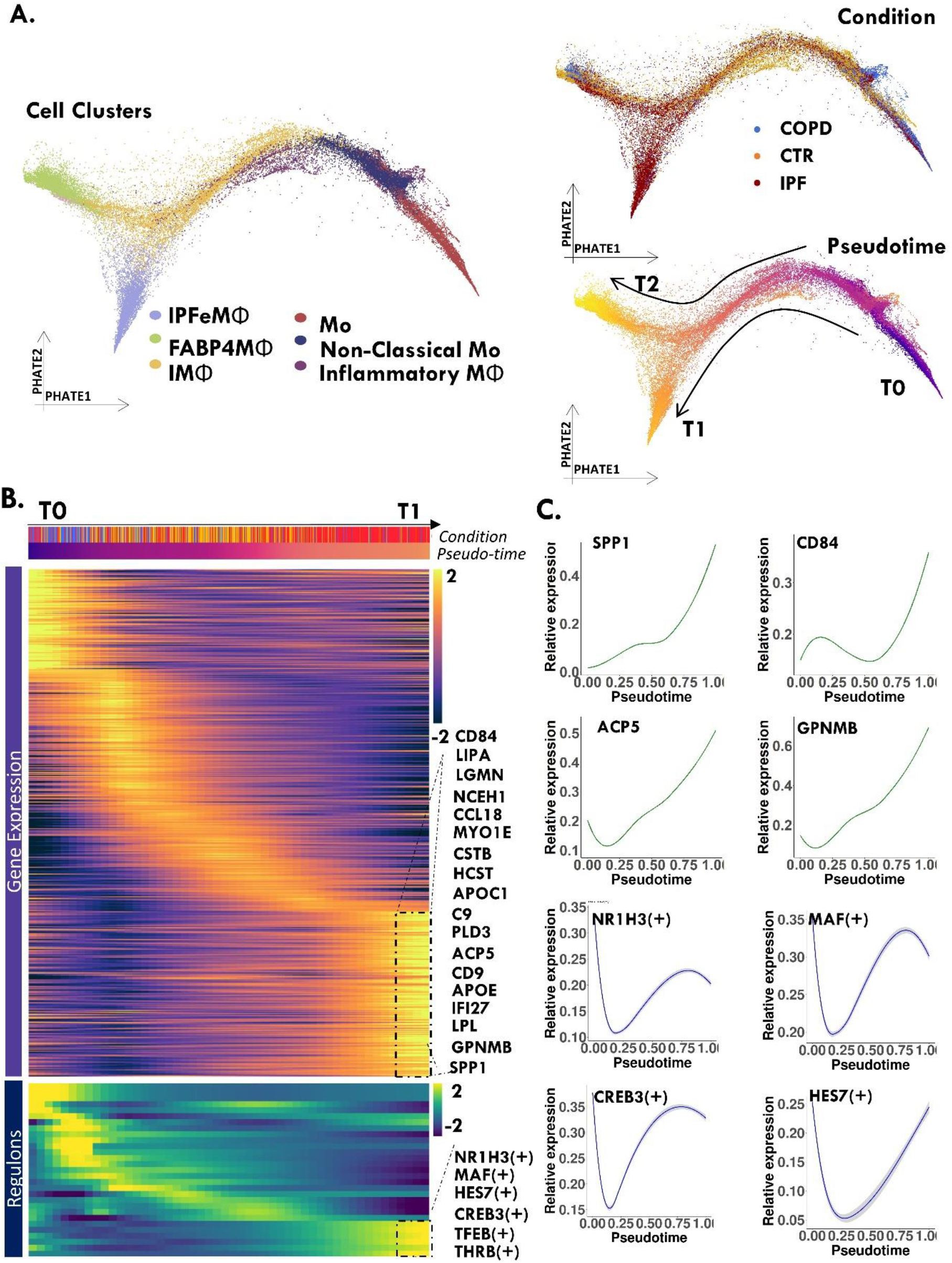
Integrated single cell RNAseq of human lungs identifies a unique macrophage population in the lungs of patients with pulmonary fibrosis. Single-cell RNA-Seq was performed on single-cell suspensions generated from lung biopsies of 29 control, 18 COPD and 32 IPF patients. **A.** PHATE embeddings of macrophage and monocyte subpopulations identifies a discrete sub-branch structure (Trajectory 1 [T1]) represented by macrophages with IPF-sample origin (IPFeM**Φ**), and a second distal branching structure (Trajectory 2 [T2]) represented by FABP4+M**Φ**. Plots are labeled by unsupervised clusters, condition and pseudotime distance. **B**. Heatmap of differentially expressed genes as a function of pseudotime distance. Top genes from IPFeM**Φ** are shown. **C**. The relative expression of select genes and regulons (X-axis) by pseudotime (Y-axis).

Genes with differential patterns of expression along their trajectory path are shown in the gene expression heatmap (Figure 1B). We observed a gradual increase in the expression of extracellular matrix (ECM) remodeling genes along the pseudotime T1 trajectory. These genes include metalloproteinases (MMP7, MMP9), secreted mediators (CCL18, SPP1, CHI3L1), proteases (CTSB, CTSK), modulators of tissue remodeling (TGBI, VIM), and enzymes (CHIT1) implicated in the pathobiology of IPF(25–29). Interestingly, a significant number of genes related to lipid metabolism were found to be differentially expressed as a function of pseudotime by IPFeM**Φ** (ex: LPL, LIPA, NCEH1, CD36), compared to FABP4+ MΦ (ex: PPARG, FABP4, FABP5), suggesting a role for differential lipid metabolism in the delineation of these two macrophage subsets.

pySCENIC (24) identified differentially expressed regulons across the T1 trajectory (Figure 1B and 1C), including a group of transcription factors differentially expressed in IPFeM**Φ** that provides insight into the potential function of this macrophage subset. NR1H3, known as Liver X receptor alpha (LXRa), is a nuclear receptor activator of SREBP-1c, with downstream effects associated with lipogenesis, cholesterol efflux, and positive regulation of M2 related genes (30)(31). MAF is a trans-activator of SPP1, which has been described in bone marrow fibrosis (32), and CREB3 and CREB3L2 are associated with ER and Golgi stress and hepatic fibrosis (33).

Triwise(34) R package identified overlapping and unique genes between the three phenotypes of the Monocyte/Macrophage trajectory path – IPFeMΦ, FABP4+M**Φ**, and monocytes. CD14 and APOE, both markers of Monocyte-derived alveolar macrophages (35), were highly specific for IPFeMΦ (Figure 2A-B). SDC2, a gene described to be expressed in alveolar macrophages in IPF (36), was also selectively expressed by IPFeMΦ. CCL18, a serum biomarker that predicts mortality (37), was mostly expressed by the IPFeMΦ phenotype. LSAMP, a neuronal membrane-bound protein, and SVIL, supervillain cytoskeletal protein, were two of the most selectively expressed genes by FABP4+MΦ (Figure 2A).

**Figure 2:**
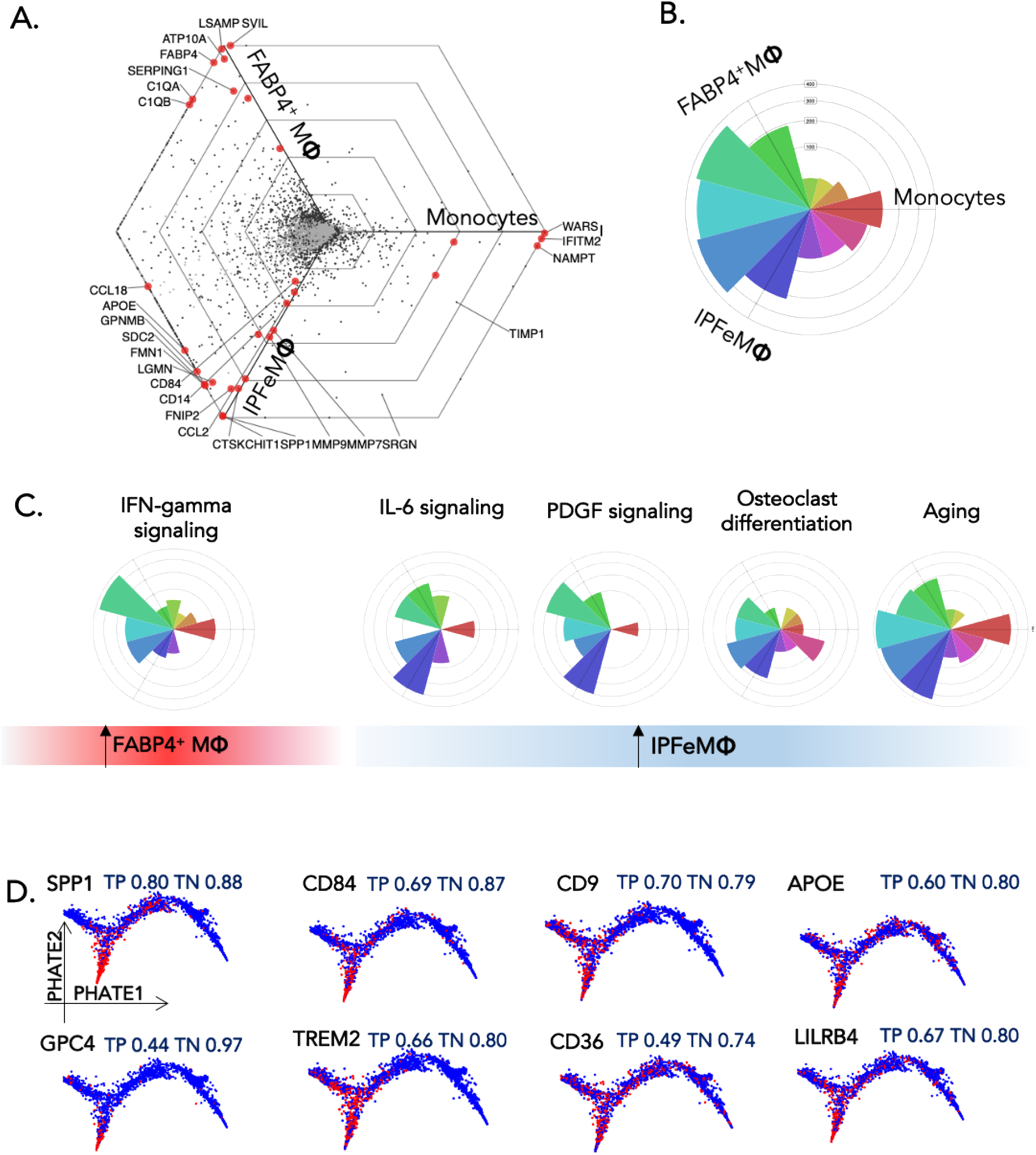
Radar plot representation of differential expressed genes between IPFeM, FABP4M and monocytes. **A.** Triwise radar plots depicting gene ontogeny (GO) enrichment analysis. The radar hexagonal plot utilizes barycentric coordinate transformation of the gene expression matrix with degree and distance from origin as outputs for each gene. Each gene is represented by a single point. The direction of a point indicates the condition in which the gene is upregulated, where 0° (Monocytes), 120° (IPFeM**Φ**), 240° (FABP4+M**Φ**), serves as landmarks. The distance from the origin, depicted by hexagonal gridlines, represents the strength of upregulation in log 2-fold-change compared to the other to comparisons Genes with the same expression in all three samples will lie in the center of the plot, regardless of their absolute expression values. B. Rose Plot, or histogram plot, shows the percentage of genes in each orientation C. Gene Ontology enrichment visualized by polar histogram plot for each fate pathway. Each color bar represents the number of genes that lie in specific degrees of the plot. The Blue bar identifies genes expressed in the 120° angle (IPFeM**Φ** specific genes), the red bar corresponds to genes in the 0° angle (monocytes), and the green bar identifies genes in the 240° angle (FABP4+M**Φ).** All other color bars are intermediate ranges between the above mention landmarks and phenotypes. Upregulation of IFN-gamma signaling was found to be a differentially expressed pathway in FABP4+ macrophages, compared to IPF expanded macrophages, which were associated with increased IL-6 signaling, PDGF signaling, osteoclast differentiation, and aging pathways. D. PHATE plots depicting the top 10 cell surface-protein coding genes ranked by minimal hypergeometric test (mHG), a non-parametric test implemented in *COMETSC* output. True positive (TP), and True negative (TN) values reported by *COMETSC* analysis are showed above each PHATE plot.

Gene ontology (GO) (38,39) enrichment analysis was implemented to identify biological processes overrepresented in these cell populations (Figure 2C and Supplementary Figure 2). The IL-6 signaling pathway was overrepresented in IPFeMΦ, with IL6 signal transducer and IL6-receptor, among the genes differently expressed in IPFeMΦ. This is consistent with prior descriptions of increased IL-6 levels in IPF, and the role of this pathway in hyper-M2 polarization (40)(41)(42). Response to Interferon-gamma, previously associated with anti-fibrotic properties(43), was overrepresented by FABP4+M**Φ.** Overall, this enrichment analysis demonstrates that immune responses are uniquely regulated between the two macrophage phenotypes.

### A unique set of cell surface markers are expressed by IPF-expanded macrophages

Using the Combinatorial Marker Detection from Single Cell Transcriptomic Data (*COMETSC*) package (44), we were able to identify the protein-coding genes for cell surface markers that could discriminate the IPFeMΦ subpopulation from other myeloid cells (Figure 2D and supplementary figure 4). SPP1, GPC4, CD84, TREM2, CD9, CD36, APOE, LILRB4, CD276 and SLAMF7, were the top 10 rank genes based on the highest minimal hypergeometric test (mHG), for the discrimination of IPFeMΦ from other macrophage and monocyte populations.

### IPFeMΦ are a mixture of alveolar and interstitial macrophages with an M2-like phenotype

Polarity of monocytes and macrophages was assessed with the two-index tool, *MacSpectrum* (45), which derives a macrophage polarization score (MPI) and activation-induced macrophage differentiation index (AMDI) for each cell. We found that IPF-expanded macrophages have a mean MPI of −2 (95% CI (−6)−3), suggestive of M2 polarity, and a mean AMDI of −13 (−95% CI (-21-3), which is associated with an immature phenotype that is characteristic of cells that originate from circulating monocytes(45) (Figure 3A).

**Figure 3:**
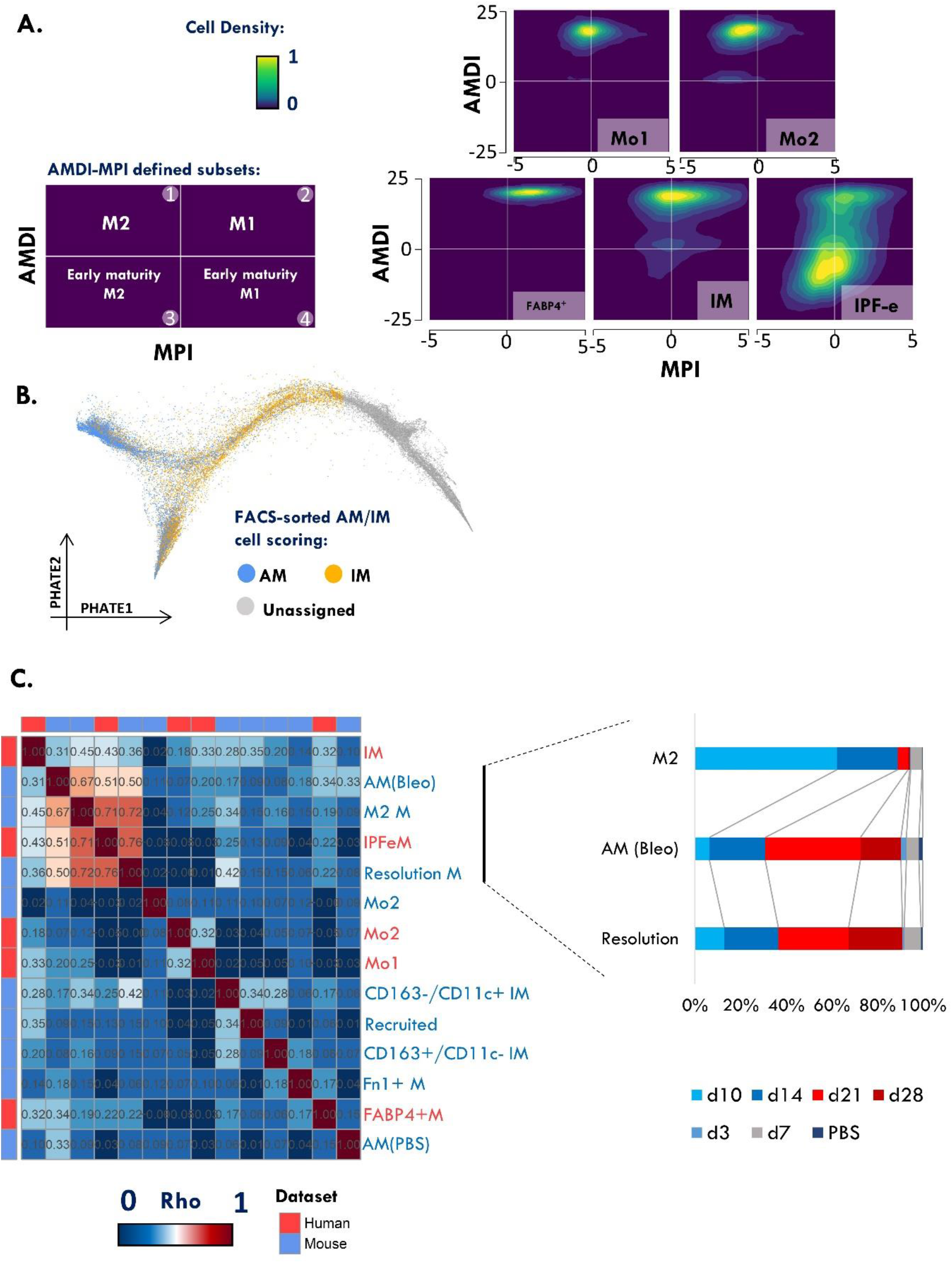
Ontogeny, maturation and tissue location of IPFeMΦ. **A.** Macrophage polarity identified by MacSpectrum (45) indices: Macrophage Polarization Index (MPI, describing M1 and M2 polarization states) on the x-axis, and Activation Maturation derived index (AMDI, describing the degree of macrophage terminal differentiation) on the y-axis. Four subsets can be identified by the relation between AMDI and MDI: Upper left quadrant (negative MPI, positive AMDI), labeled ‘M2-like macrophages’; Lower left quadrant (negative MPI, negative AMDI), labeled as ‘M2-like Pre-activation cells’; Upper right quadrant (positive MPI, positive AMDI), labeled as ‘M1-like macrophages’; Lower right quadrant (positive MPI, negative AMDI), labeled as ‘M1-like Pre-activation cells’. Utilizing this matrix, the polarity of monocyte (Mo) and macrophage subsets were identified. **B.** An external data of FACS-sorted bulk was used to score cells based on their similarity to either FACS-sorted IM or AM cells. Cells were labeled as AM or IM if the score difference for either category is higher than 0.05. IPFeMΦ demonstrated both IM and AM characteristics. **C.** Correlation matrix of macrophage/monocyte cell subpopulations between our dataset (red) and a bleomycin mouse model (blue), Matrix cells are colored by Spearman Rho correlation coefficient. Hierarchical clustering was implemented to order the clusters. *Right inset*, showing distribution of macrophages from the Bleomycin model split by day after bleomycin administration.

To identify the ontogeny of the IPFeM**Φ**, published bulk RNA-seq datasets (46) from FACS-sorted IM and AM were used to construct distinct signatures for each macrophage subset. We calculated similarity scores for each cell as previously described (47), and determined the correlation coefficient of each cell to the differentially expressed genes in each of the FACS-sorted IM/AM signatures, allowing each cell to be labeled as IM or AM (Figure 3B). Interestingly, IPFeM**Φ** comprised a population of cells categorized as both IM (51%) and AM (49%), spanning the beginning (IM) and the terminal tip (AM) of the sub-branch structure. This suggests a continuum of differentiation, where IM is an intermediate cell-state and AM is the terminal phenotype. On the other hand, FABP4+ macrophages were predominantly scored as AM (83%).

To understand the temporal emergence of IPFeM**Φ**, we integrated our data with the publicly available dataset derived from a bleomycin mouse model of pulmonary fibrosis(47). We performed correlation analysis of the myeloid compartment from this external dataset using their original cell labels. IPFeMΦ showed the highest correlation coefficient with mouse cells labeled originally as ‘M2 macrophages’, ‘AM(Bleo)’ and ‘resolution macrophages’. These populations were represented in samples from day 10 and 14 (M2 macrophages) and day 21 and 28 (AM(Bleo), Resolution Macrophages). These four groups-IPFeMΦ, M2 macrophages, AM(Bleo) and resolution macrophages, showed the highest correlation coefficients from the entire comparison, suggesting that IPFeM represent a population of transitional pro-fibrotic macrophages, that expands after lung injury and remain until late stage of fibrotic response (Figure 3C). As an external validation, we utilized the available human lung dataset from Habermann et al. (48), comprised of 10 PF and 20 controls. We were able to identify a population of macrophages that shared a similar gene expression and are predominantly found in fibrotic lungs (Supplementary Figure 5).

### IPFeMΦ exhibit unique cell-cell interactions with cells in the fibrotic niche

To better understand the cellular interactions between IPF macrophages and other cell types, a ligand-receptor (L-R) and ligand-target (L-T) interaction map was built using *NicheNet* (34) R package. IPFeMΦ were selected as sender cells and myofibroblasts, vascular endothelial (VE) cells, and aberrant basaloid epithelial cells were selected as receivers. A circular plot with the most significant L-T interactions is shown in Figure 4A, and specific L-R and L-T interactions are shown in plots in Figure 4B. An alternative analysis, utilizing IPFeMΦ as receivers, is presented in Supplemental Figure 6. We identified specific IPFeMΦ ligands that interact with myofibroblast intracellular targets and inferred receptors. Prioritized potential ligands in IPFeMΦ included TGFB1, TNFSF13B, SPP1, GPNMB, and PIK3CB. Intracellular targets in Myofibroblasts that interact with the prioritized ligands were multiple Collagen related genes (COL1A1, COL1A2, COL1A3), ECM-modulators (MMP2, TIMP1 and TIMP3) and ECM-components such as glycoprotein VCAN, Elastin (ELN), FN1, FBN1, and PALLD. Inferred receptors in myofibroblasts that interact with TGF-β were TGBR1, TGFBR2, SDC2, Vitamin D receptor (VDR), ACVRL1, and BMPR1A (Figure 4B).

**Figure 4:**
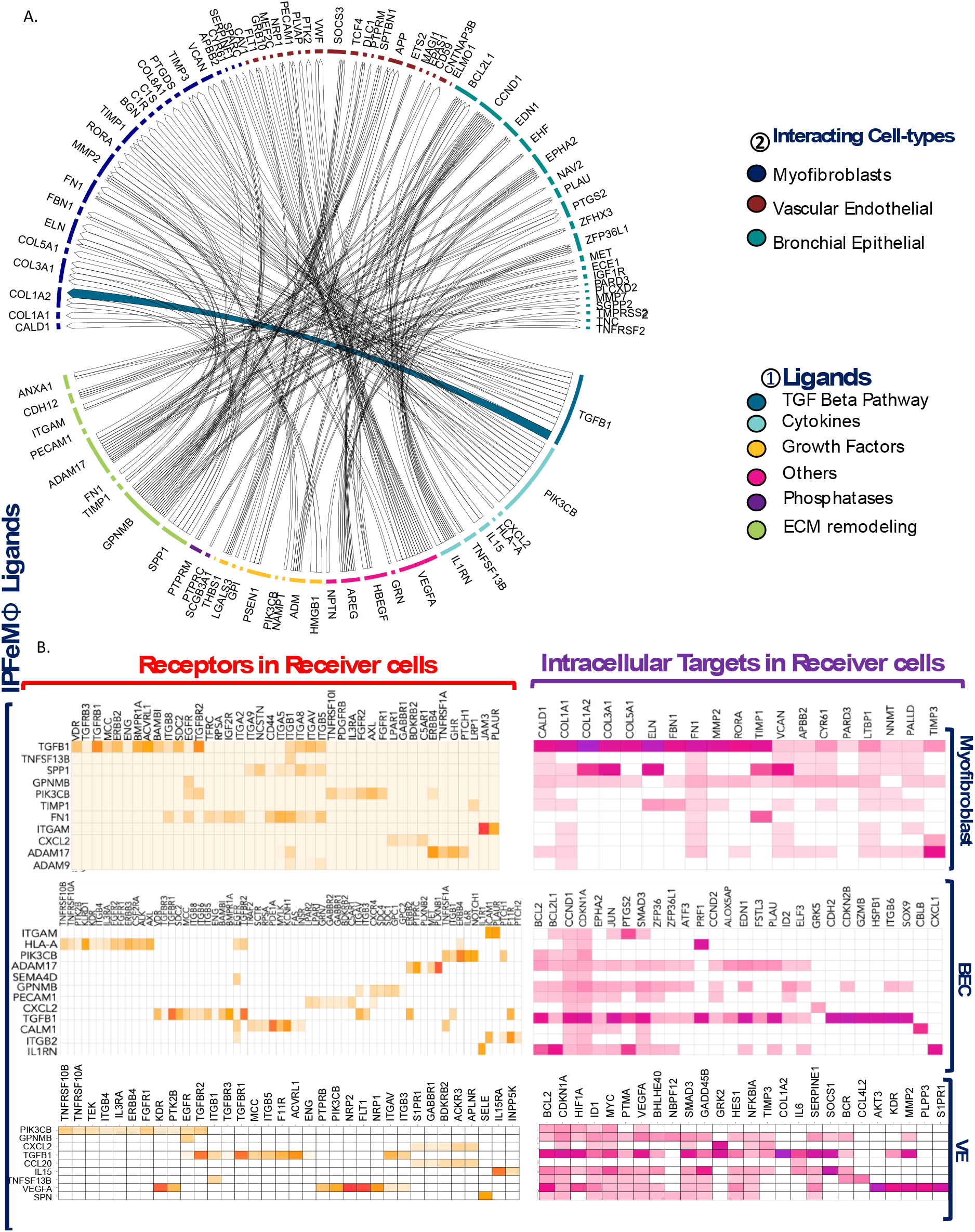
IPFeMΦ interacts with multiple cell types in the fibrotic niche, with potential activation of pro-fibrotic response and cell survival in different structural cells. **(A)** Circle plot showing links between **(1)** predicted ligands from IPFeM**Φ**, and **(2)** intracellular targets in receiver cells – myofibroblasts, aberrant basaloid epithelial cells (BEC) and vascular endothelial (VE) cells. Colors were assigned by the biological nature of the interaction: TGF-B pathway, Cytokines, Growth Factors and ECM related genes. Targets were color-coded based on the cell-type that gene represents. **(B)** Schematic representation of the *NicheNet* analysis of upstream ligand-receptor pairs. **(C)** Potential receptors expressed by receiver cells – myofibroblasts, BEC, and VE cells-associated with each potential ligand expressed by IPFeM**Φ**. Potential intracellular targets in different receiver cells that interact with the upstream ligands from IPFeM**Φ**.

The ligands that were prioritized in the IPFeMΦ-Aberrant Basaloid cell interaction included ITGAM, HLA-A, ADAM17, SEMA4D, TGFB1, among others. Of these, TGFB1 was the ligand with the greatest number of intracellular targets, interacting with genes related to cell survival (CCND1, CCND2, BLC2, BCL2L1), cell senescence (CDKN1A, CDKN2B), Ephrin pathway (EPHA2, with knockout of this pathway described as protective in Bleomycin models (49)), Endothelin-1 (EDN1), Urokinase (PLAU), and SOX9, a known regulon overrepresented in these aberrant epithelial cells (18) and other forms of lung injury (50).

Interactions between VE cells and IPFeMΦ exhibited similar prioritized ligands to the above-mentioned cells, but slightly different Intracellular targets are potentially activated. TGFB1 connected with genes related to cell survival (BCL2, CDKN1A, HIF1A), endothelial progenitor transcription factors (ID1, MYC), cytokine production (IL6), and inhibition of fibrinolysis (SERPINE1). VEGFA ligand in IPFeMΦ interacted with molecules related to angiogenesis (AKT3, KDR, S1PR1) and cytokine signaling (PLPR3). IL-15 – an IPFeMΦ ligand, interacted with multiple targets, including cytokine signaling (SOCS1 and IL6), growth arrest transcription factors (GADD45B/MyD118, HES1, BHLHE40), and ECM-remodeling (TIMP3, MMP2) (Figure 4B).

Overall, we used *NicheNet* to predict the primary ligands utilized by IPFeMΦ to interact with the fibrotic niche, and the potential downstream effects of these interactions in select receiver cells. From our analyses, IPFeMΦ ligands may act to increase collagen and ECM-related protein production in myofibroblasts. In basaloid epithelial cells, IPFeMΦ ligands may regulate activation of epithelial-mesenchymal transition and cell survival through initiation of senescence and activation of fibrinolysis. In vascular endothelial cells, IPFeMΦ interaction may augment expression of inflammatory molecules, ECM-remodeling proteins, fibrinolysis inhibitors and molecules associated with cell-growth arrest.

### Utilizing CyTOF to validate the presence of an expanded macrophage population in IPF patients

To identify and confirm the existence of the expanded macrophage population in IPF compared to COPD and control lung tissues, we stained singled cell suspensions prepared from lung tissues with a CyTOF antibody panel containing 35 cell-surface markers that included classical macrophage markers and novel surface markers as predicted by scRNAseq (Supplementary Table 1). The data was analyzed using Cytobank (https://cytobank.org) and as previously described by Ng. et al(51). We first manually gated for common leukocyte populations within the intact, singlet, liveCD45+ cell subset to determine the proportion of cell populations found between the three groups (Figure 5A). We then applied the t-distributed stochastic neighbor embedding (t-SNE) dimensionality reduction on the dataset to visualzie the cell populations (Figure 5B). Using X-shift algorithm(52) for k-nearest neighbor estimation, we identified the optimal number of clusters from the CD45+ gate, which we found to be 30. The SPADE algorithm (53) was subsequently used to identify clusters of phenotypically similar cells. The algorithm revealed one cluster, cluster 25, to be more abundant in lungs from IPF patients compared to healthy and COPD controls (Figure 5D). A heatmap of the mean expression levels of the phenotypic markers in each cluster (Figure 5C) revealed that cluster 25 is comprised of a hybrid/transitional population that has both alveolar and interstitial macrophage markers (HLADR+, CD11b+, CD206+). Notably, there was strong expression for CD84, CD36 and CD64 in cluster 25 (Figure 5G-I), confirming some of the unique surface markers predicted by scRNAseq. Overall, the data confirmed several novel markers in a unique macrophage population that is expanded in IPF.

**Figure 5:**
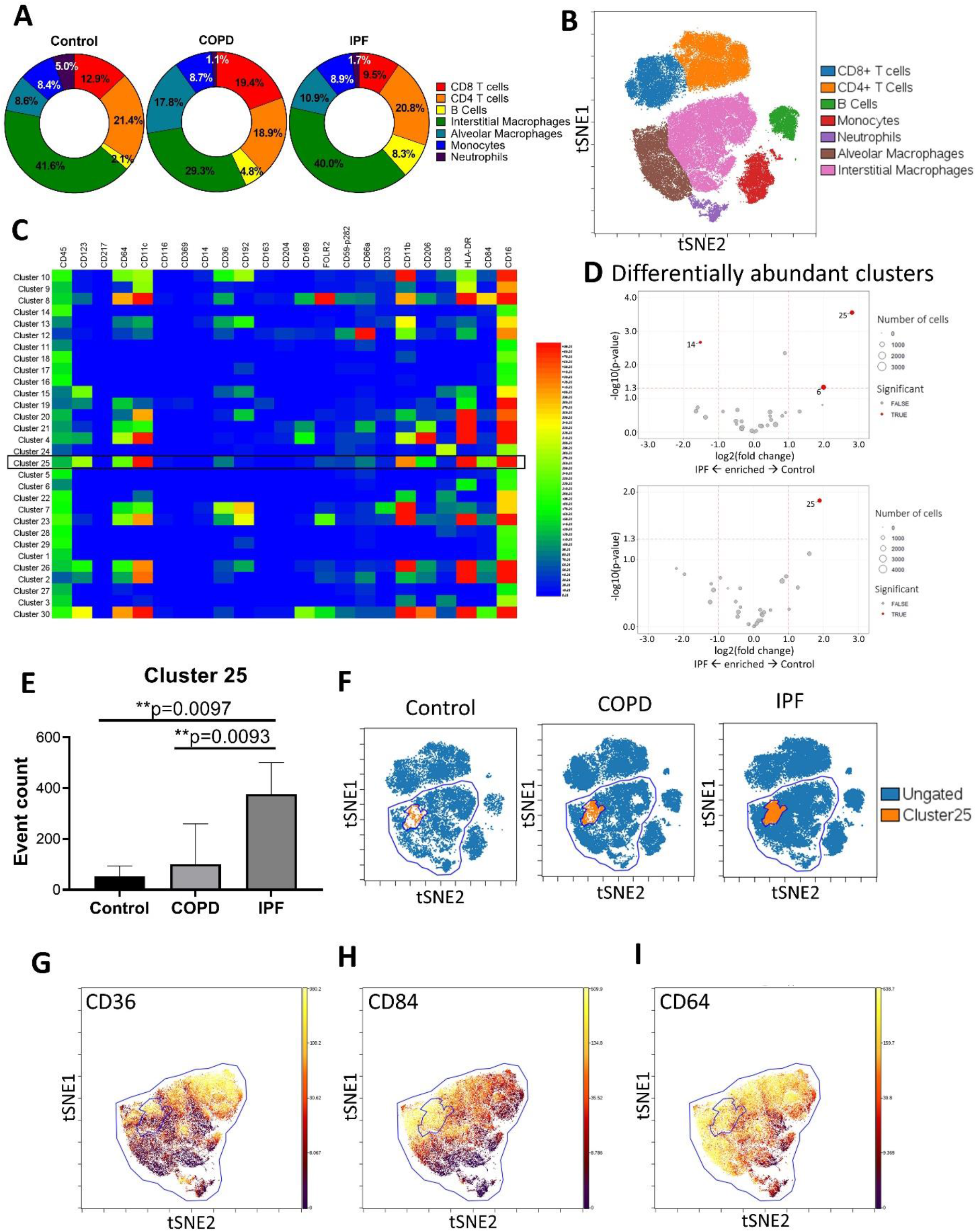
CyTOF profiling of the human lung reveals a uniquely expanded macrophage cluster in IPF compared to COPD and control subjects. Human single cell tissue digests were thawed, enriched for viable cells, stained with metal-conjugated antibodies, and processed for CyTOF profiling and analysis. **(A)** Cell abundance expressed as percentages in the Donut Chart and **(B)** visualized using a t-SNE plot. **(C)** Heatmap showing the mean expression of all markers following the determination of optimal number of clusters using k-nearest neighbor estimation. **(D)** Volcano plots of clusters identified by SPADE analysis (Control vs. IPF and COPD vs. IPF) depicting differentially abundant clusters. (**E**) The relative abundance of the expanded cluster (cluster 25, orange) in IPF **(F)** t-SNE plots illustrating the expanded macrophage cluster and expression of **(G)** CD36, **(H)** CD84 and **(I)** CD64 in that cluster. The color gradient indicates the marker expression intensity.

### Cell surface expression of CD84++ and CD36++ distinguishes the hybrid macrophage population expanded in IPF patients

Our scRNAseq and CyTOF analyses validated the presence of an expanded macrophage population that may have pathologic importance for chronic fibrotic lung disease. To better understand the biological heterogeneity of CD84 and CD36, and whether their expression levels delineate subpopulations that distinguishes disease states, we performed flow cytometric analysis on 10 control, 10 IPF and 10 COPD lung digests and performed a gating strategy similar to that described in Bharat *et. al* (54) (Figure 6A). In addition to anti-CD36 and anti-CD84, we included a viability marker (Zombie dye), anti-CD45, CD15, HLA-DR, CD206 and CD169, to characterize the live lung macrophage population (Figure 6A). We used CD206 and CD169 to delineate alveolar (CD206+CD169+) and interstitial (CD206+CD169-) subsets, and to determine the relative expression levels of CD84 and CD36 in these macrophage subsets. We identified a CD84++ and CD36++ macrophage population that is uniquely increased in the lungs of IPF compared to control and COPD subjects (Figure 6B-E). Of note, there was little to no increase in the CD84+ (low) and CD36+ (low) macrophage population in the IPF disease cohort (Figure 6F-I). In contrast, there was marked reduction either in the proportion or the mean fluorescent intensity (MFI) of CD84+ (low) and CD36+ (low) macrophage population in COPD (Figure 6F-I). These findings validate the CyTOF data and further confirms the presence of an expanded population of CD84++CD36++ macrophages in the lungs of patients with IPF.

**Figure 6:**
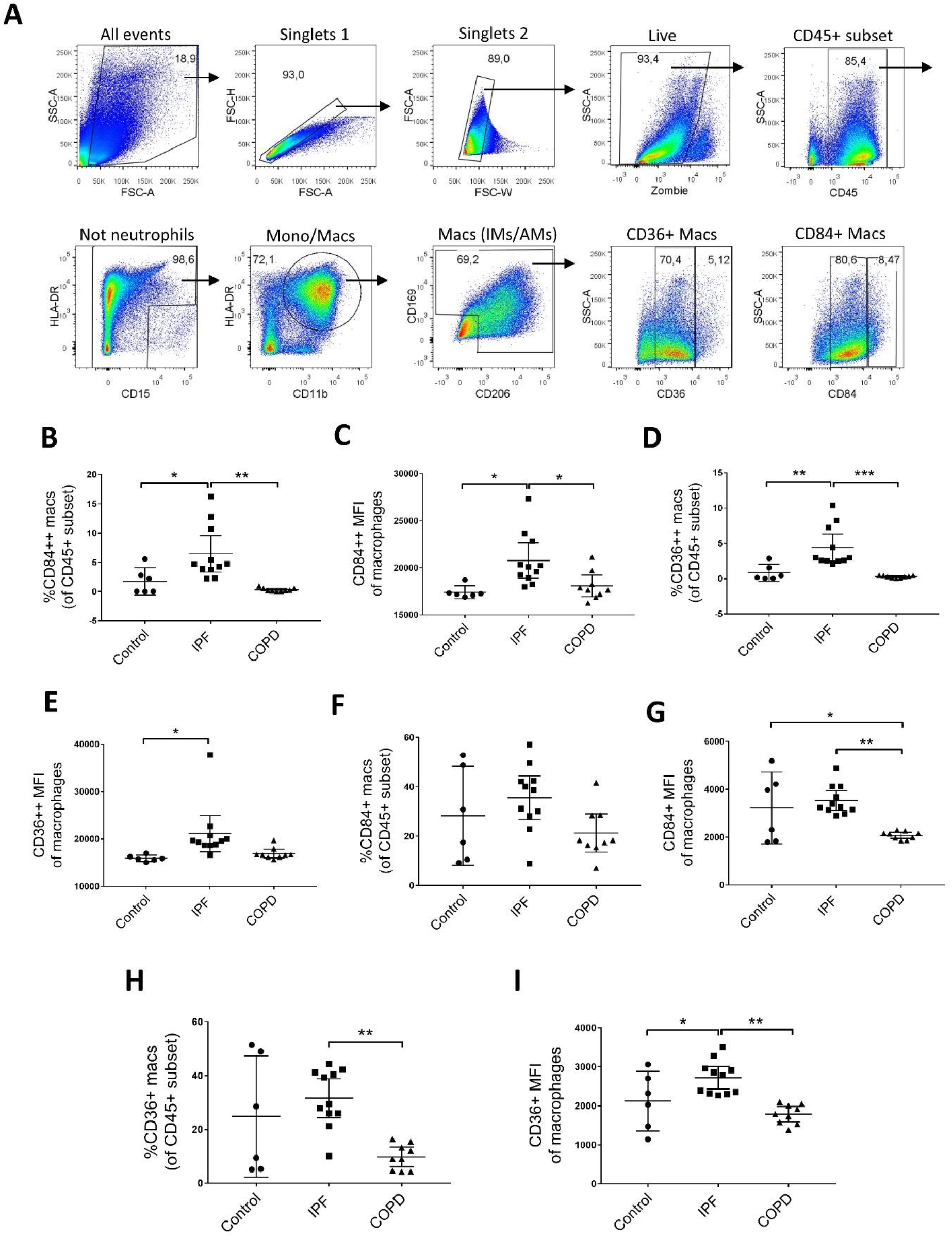
Flow cytometric analysis of human lungs validates the presence of an IPF-expanded macrophage population. Biobanked human lung tissue digests were thawed and stained with viability marker, antibodies to delineate the macrophage populations, and scRNAseq-predicted surface antibodies against the expanded IPF subset. **(A)** A gating strategy showing detailed sequence of flow plots leading to the interstitial and alveolar macrophage compartment. Subset of immune cells were gated as follows: Neutrophils (CD45^+^HLADR^-^), Macrophages (CD45^+^HLADR^+^CD15^-^CD11b^+^CD206^+^CD169^+^), CD36^+^/ CD36^++^ macrophages and CD84^+^/ CD84^++^ macrophages. The relative proportion of **(B)** CD84^++^, **(D)** CD36^++^, **(F)** CD84^+^, and **(H)** CD36^+^ macrophages in the CD45^+^ subset, and the corresponding mean fluorescent intensity of **(C, G)** CD84 and **(E, I)** CD36. Data were analyzed with one-way ANOVA using Tukey’s multiple comparisons test. *, p < 0.05; **, p < 0.05; ***, p < 0.05.

### Utilizing the L100CDS2 dataset to predict drugs that reverse the transcriptional signature of IPF-specific macrophages

We used the publicly available L1000 Characteristic Direction Signature Search Engine (L100CDS2) to predict drugs that could reverse the signature of IPF-specific macrophages and then tested several candidate molecules *in vitro* using a human-derived macrophage system and CCL18 secretion – an IPF specific biomarker(55)- as the primary endpoint. We used L1000 Fireworks display (L1000FWD) (56)(57) - a t-SNE reduction of the multiple drug signatures into two-dimensions, to localize the drugs of interest (Figure 7A-B). The identified candidate drugs that could potentially reverse the profibrotic signature were then categorized by their mechanism of action on specific biological processes (ex: STAT3/ HSP90/ proteasome/ fatty acid synthase/ MMP2 inhibition and apoptosis induction). We decided to focus our attention on STAT3 modulators due to: 1. the presence of multiple pharmacological STAT3 regulators among the top pharmacological compounds, and 2. the implication of STAT3 signaling in pro-fibrotic and M2-like macrophage responses(41) (58). We selected three drugs: cinobufagin [Cino], cucurbitacin I [Cuc], and cryptotanshinone [CRT], and tested their ability to inhibit the pro-fibrotic M2-like macrophage phenotype, including hyper M2-like macrophages preconditioned with IL-6, as previously described (40)(41)(42). We performed *in vitro* viability testing using multiple drug concentrations on control, M2-like and hyper M2-like macrophages. A range of concentrations with 10-fold differences were tested to exclude the toxic drug concentrations that reduced cell viability, as demonstrated by the MTS viability assay (Figure 7C). Next, we performed functional assays by testing suitable drug concentrations on CCL18 production by M2-like macrophages and TNFα production by pro-inflammatory M1-like macrophage phenotypes. As expected, IL-6 acted synergistically with IL-4 and IL-13 to hyper polarize macrophages towards the M2 phenotype by increasing CCL18 production (Figure 7C). At 100nM, both Cuc and CRT reduced IL-6-mediated CCL18 production without impacting TNFα production from pro-inflammatory macrophages (Figure 7D). Altogether, these results demonstrate the feasibility of predicting potential drug targets using scRNAseq signatures, which can subsequently be confirmed using established *in vitro* assay systems.

**Figure 7.**
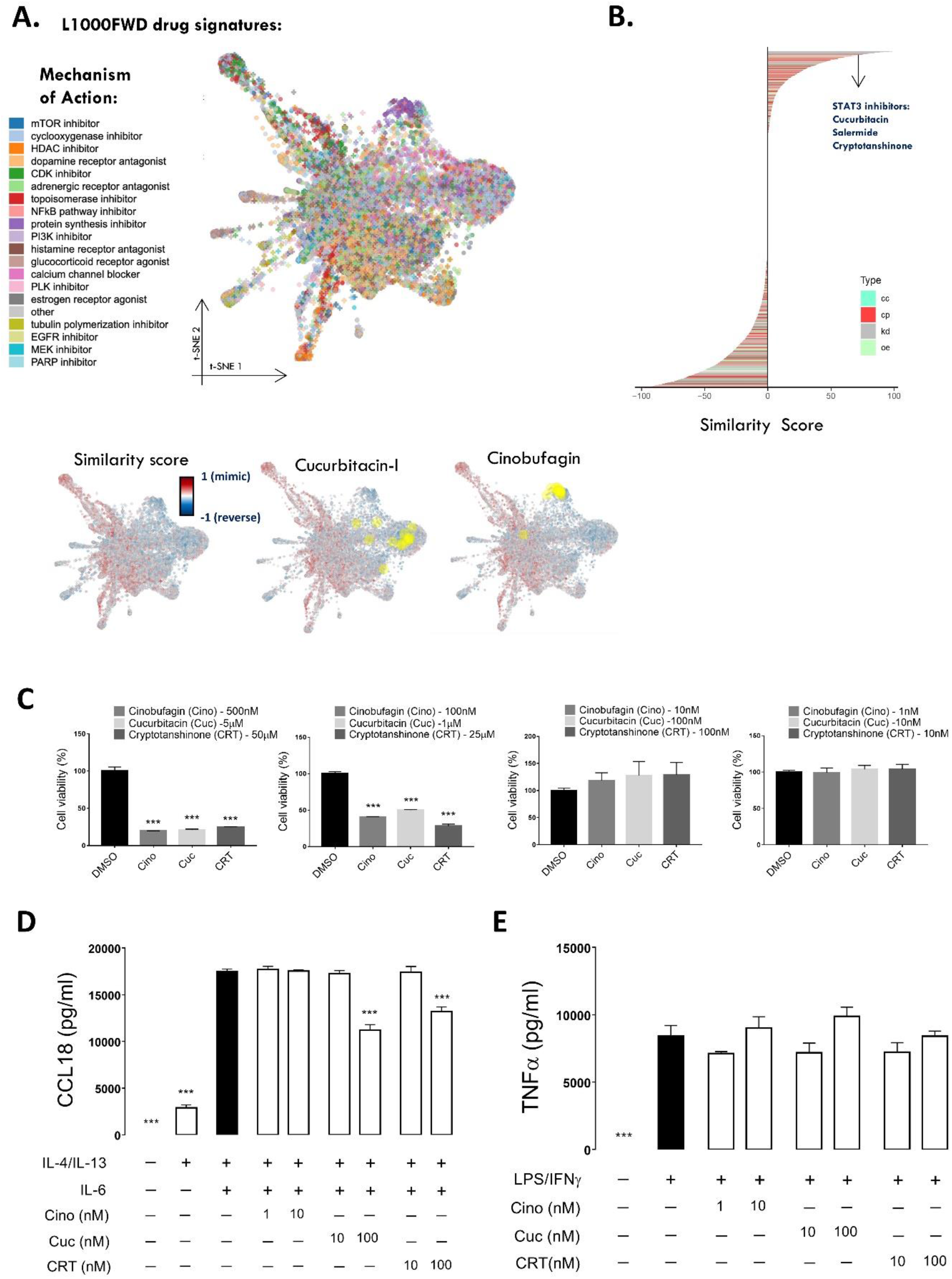
Prediction and validation of drugs that reverse the transcriptional signature of IPF-specific macrophages. LINCS L1000 characteristic direction signature search engine (L1000CDS2) algorithm was used to predict drugs that reverse the transcriptional signature of IPF-specific macrophages. Functional ability of selected drugs was tested in a CCL18-producing THP1 derived macrophage system. **(A)** L1000 Firework Display (L1000FWD) t-SNE visualization tool was used to identify drug signatures and their corresponding mechanisms of action (MOA) that could reverse the IPFeMΦ signature. **(B)** Perturbation signatures were ranked based on the similarity score, revealing multiple drugs that interact with STAT3 pathway-Cucurcitabin, salermide, cinobufagin, and cryptotanshinone. Cucurcitabin and cinobufagin were represented and highlighted on a L1000FWD t-SNE plot. **(C)** *In vitro* toxicity/cellular viability of THP1-derived macrophages upon treatment with Cinobufagin, Cucurbitacin I and Crypotanshinone. Concentrations of secreted **(D)** CCL18 and **(E)** TNFα from THP1-derived macrophages polarized with IL-4/IL-13/IL-6 and LPS/INFγ. Data were analyzed with one-way ANOVA using Tukey’s multiple comparisons test. ***, p < 0.05; where *** represent a difference between the indicated condition and the reference control (black bars).

To investigate whether the predicted drugs reverse the phenotype of IPF macrophages in a disease relevant setting, we utilized an IPF precision cut lung slice (PCLS) system as an *ex-vivo* translational tool to modulate IPF macrophage biology (59)(60)(61). These slices were cultured fresh and demonstrated a great degree of viability (90%) over the course of 4 days based on flow cytometric live/dead analysis (Supplementary Figure 7). We gated for the target populations using fluorescent minus one (FMO) controls to ensure a specific gating strategy. Fresh IPF PCLS were treated with Cuc and CRT (100nM each) for 48 hours and then processed for flow cytometric analysis and RNA extraction. Supernatants were saved for the assessment of secreted mediators from fibrotic tissues. Not only were the proportions of CD84+, CD36+ and CD64+ macrophages reduced in response to Cuc and CRT in the fibrotic slices (Figure 8A-C), but there was also a significant reduction in the mean expression of these markers by MFI on macrophages from PCLS (Figure 8D-F). Moreover, Cuc reduced CCL18 levels, as shown by the reduction in mRNA expression and as measured by ELISA (Figure 8G-H). These findings suggest that Cuc can modulate macrophage biology in IPF slices with a meaningful reduction in secreted CCL18.

**Figure 8.**
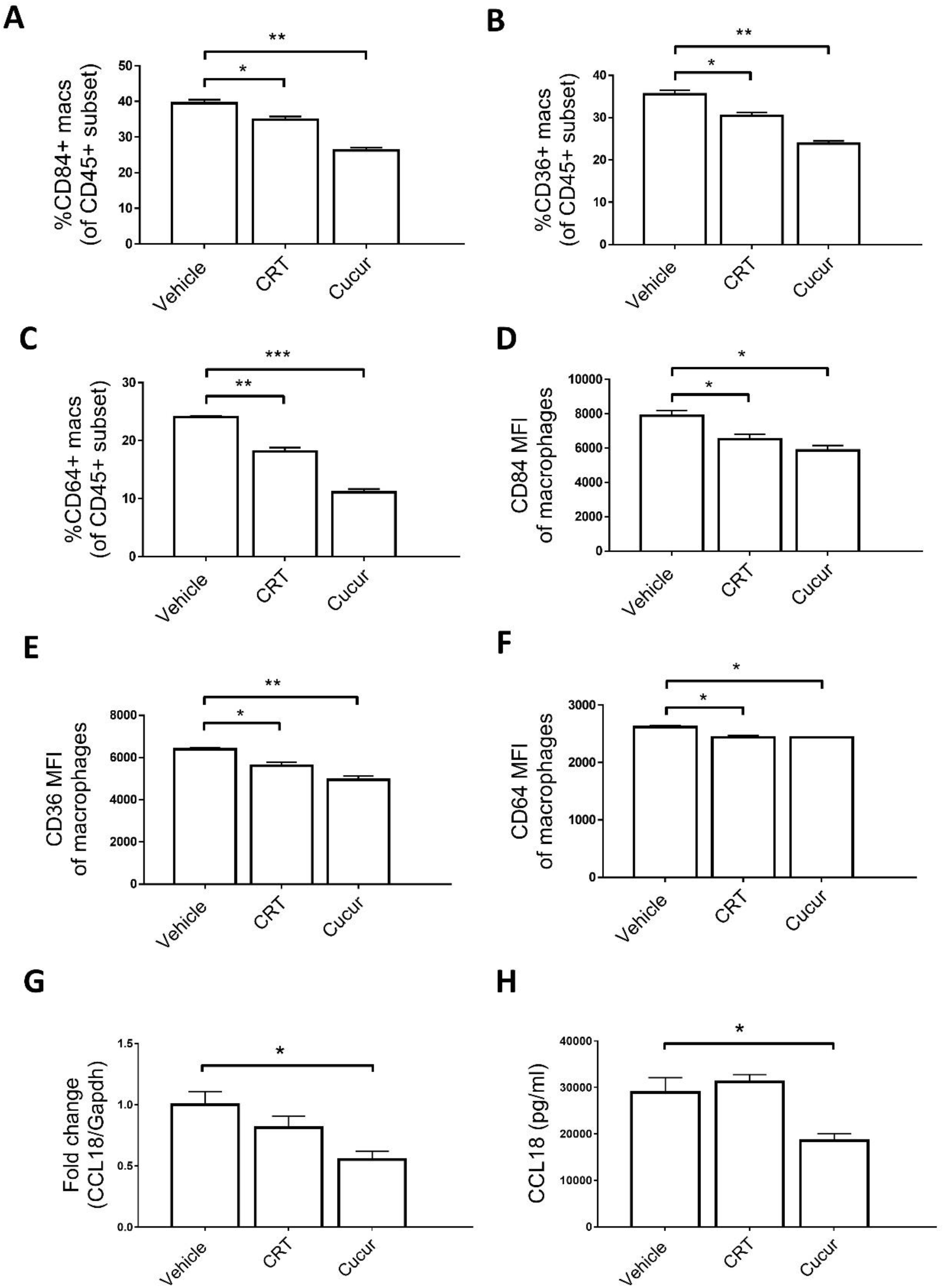
Pharmacological modulation of IPF precision cut lung slices (PCLS) reduces the accumulation of the aberrant macrophage population and CCL18 secretion. End-stage IPF lung was inflated with low melting dose agarose and slices were cultured in media. Cryptotanshinone and cucurbitacin (100nM) were then added to the cultured slices for 48hrs. After 48 hours, supernatant was collected, and tissues slices were subjected to flow cytometric analysis and RNA extraction. **(A-C)** Proportion of CD36+, CD84+ and CD64+ macrophages shown in CD45+ cells. **(D-F)** MFI of CD36, CD84 and CD64 are shown within their respective populations. **(G)** Gene and **(H)** protein expression of CCL18 as assessed by RT-PCR and ELISA, respectively. Data were analyzed with one-way ANOVA using Tukey’s multiple comparisons test. *, p < 0.05; **, p < 0.05; ***, p < 0.05.

## DISCUSSION

Despite recent advances in the understanding of macrophages during the course of IPF (16–19)(48)(62), an integrated multi-omics approach incorporating complex 3D systems are needed to precisely interrogate the existence of pro-fibrotic macrophages and to provide in-depth insights into the efficacy of targeted therapeutics. To this end, we report the first study to our knowledge of utilizing and linking various techniques (scRNAseq, CyTOF and PCLS) to understand macrophages implicated in fibrotic lung disease. Using human lung biopsies from patients with end-stage IPF and COPD, and rejected donor lungs from healthy controls, we showed that IPF lung tissues possess a uniquely expanded macrophage population compared to COPD and control lungs. By utilizing various transcriptomic approaches, we confirmed a monocyte-derived lineage origin to the expanded macrophage cluster and assigned it a predominant M2-like fate with IM and AMs features, suggesting a transitional macrophage phenotype. We identified novel cell discriminatory surface markers and undescribed transcription factors -NH1R3, MAF, HES7, CREB3, THRB-that may drive their pro-fibrotic function. Moreover, we shed some light into the potential interaction of these IPF macrophages with the fibrotic niche, particularly myofibroblasts, aberrant basaloid epithelium, and vascular endothelial cells. Using some of the surface markers predicted by *COMETSC*, we confirmed the expansion of this population in IPF lungs in an unsupervised clustering algorithm using high dimensional analysis of CyTOF data derived from lung digests. More specifically, we found that the expanded macrophage cluster expresses CD84 and CD36. Additionally, through the cellular phenotyping analysis studies, we confirmed the initial findings from scRNAseq that this cluster possesses features of both alveolar macrophages and interstitial macrophages, adding evidence to the existence of a transitional macrophage population that contributes to fibrotic disease pathology(63). Additionally, we demonstrated that we can utilize the IPF-specific macrophage signature derived from scRNAseq and the LINCs dataset to predict molecules that have the ability to reverse the IPF-specific macrophage phenotype, as demonstrated by the remarkable reduction in CCL18, a sensitive marker reflecting *in-situ* macrophage reprograming towards a pro-fibrotic phenotype(64). Taken together, our findings are consistent with the existing literature that describes distinct markers in IPF reflective of multiple subpopulations of macrophages, and that a unique subgroup of transitional macrophages is involved for the induction of pro-fibrotic response(16,19). We hypothesize that the fibrosis-expanded cluster, marked by high surface expression of CD84 and CD36, is a discrete population expanded in IPF, and may be specifically targeted in fibrosis as a novel therapeutic approach (please see figure 9 for a visual summary highlighting the study design).

**Figure 9.**
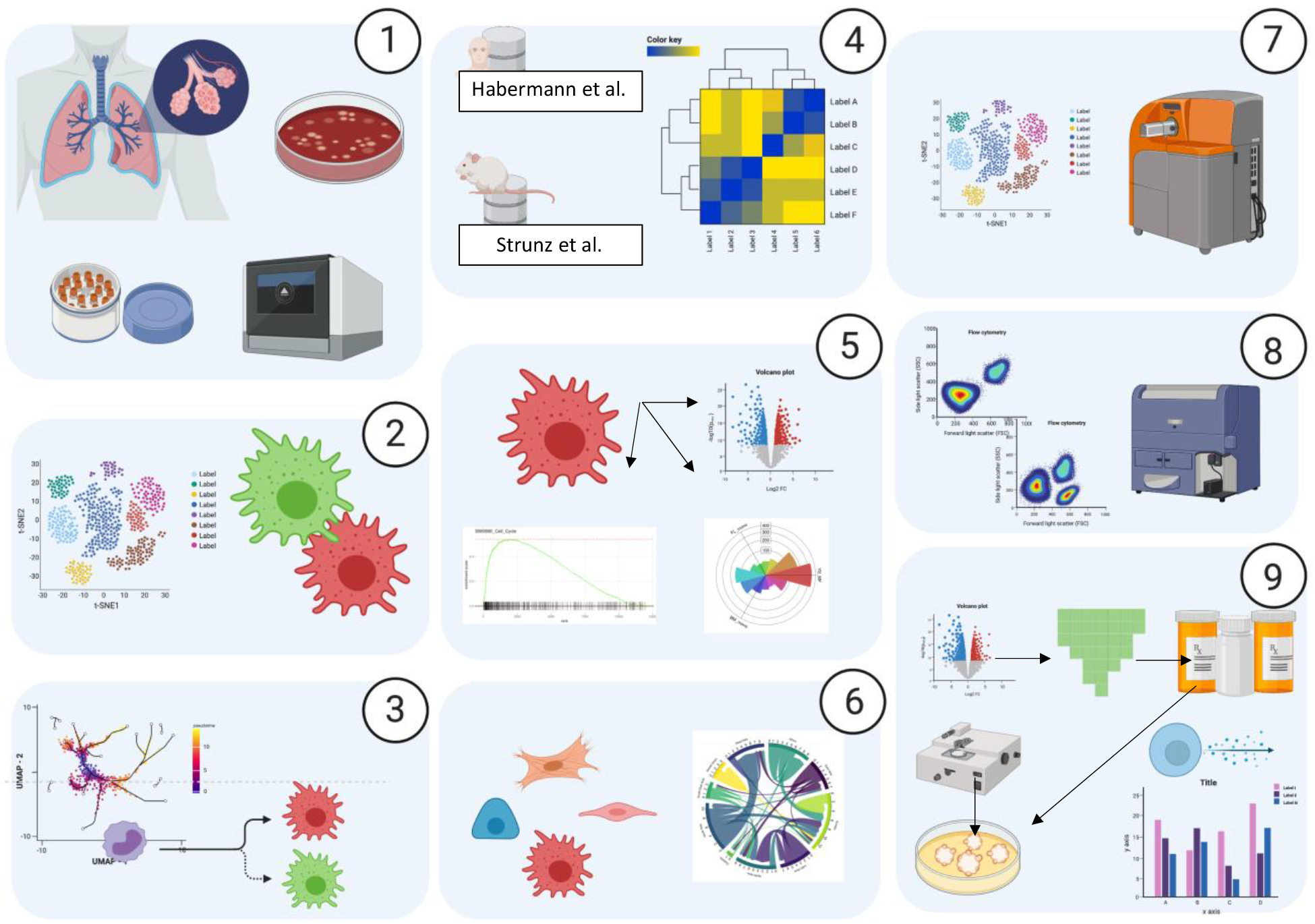
General Overview of the study design. **1.** Tissue procurement and dissociation into single cell suspensions that were cryopreserved and alter thawed and passed throught 10x pipeline. **2.** Digitalized cells were processed, myeloid compartment identified, and subsets of macrophages were characterized. **3.** Pseudotime and trajectory inference analysis of monocyte/macrophages. **4.** Correlation with external human and mouse datasets. **5.** Characterization of the IPFeM, DE analysis, gene set enrichment, identification of surface markers, and polarization and activation states with Macspectrum. **6.** Cell-cell interactions, with IPFeM as sender. **7.** CyTOF of myeloid compartment confirming findings of scRNAseq 8. FC confirmation of surface markers. **8.**. **9.** Drug molecules inferred from gene signatures using LINCS1000, were used on PCLS from IPF tissue, and CCL18 as endpoint was measured.

Our findings add significant clarity to the prior macrophage phenotyping studies in lung fibrosis performed using flow cytometry/scRNAseq, which showed that pro-fibrotic macrophages are a heterogenous pool of cells spanning the monocyte-to-macrophage differentiation spectrum. Initially, Misharin *et al*. found a greater degree of resemblance between the transcriptional signature of IPF macrophages and murine monocyte-derived alveolar macrophages, suggesting that monocyte-derived alveolar macrophages contribute to driving fibrotic responses(46). Subsequently, the same group extended the above finding with a scRNAseq approach demonstrating that there are distinct subsets of macrophages that differ in quantity between donor control lungs and IPF lungs(16). Our data is consistent with other findings that plasticity in the macrophage phenotype does exist in IPF(16,17)(19)(65) and that a plastic M1-/M2-like and a pro-resolving phenotype may co-exist depending on local tissue cytokines and growth factors. In addition, while it is difficult to trace the origin of macrophages (alveolar macrophages vs. interstitial macrophages) based on transcriptional approaches, our utilization of the multidimensional scRNAseq, mass cytometry and flow cytometric analyses on lung tissues further consolidated the pro-fibrotic transitional macrophage phenotype hypothesis in IPF. Moreover, our identification of CD84 and CD36 as discriminating surface markers for a hybrid alveolar macrophage/interstitial macrophage subtype in IPF provides a cellular basis to better understand their biology as a potential target for therapeutics in IPF.

Limited data exist regarding surface-expression levels of CD84 and CD36 as key novel molecules that phenotypically characterize a macrophage subset in IPF. CD84 is a surface molecule that is a member of the signaling lymphocyte activation molecule (SLAM) family of proteins. It forms homophilic dimers by self-association and has been shown to contribute to the survival of chronic lymphocytic leukemia (CLL) cells(66–68). Recently, Lewinsky *et al*. showed that CD84 expression up-regulates PDL1 expression on CLL cells and PD1 expression on T cells, which contributes to an exhausted T cell phenotype and suppression of T cell responses(69). On that note, Celada *et. al* demonstrated that IPF patients have an increase in PD1^+^CD4^+^ T cells relative to healthy controls, and these cells exhibit pro-fibrotic characteristics *in vitro* (70). By blocking PD1 in the murine bleomycin model of pulmonary fibrosis, fibrotic effects were significantly attenuated. Given this, the presence of an expanded CD84++ macrophage subset in IPF suggests that high expression of CD84 may control pro-fibrotic immune responses in the lung by regulating the expression of PDL1/PD1 on T cells or other cell types in the neighboring microenvironment, a hypothesis that warrants further investigation. Interestingly, CD36 is a macrophage scavenger receptor that mediates a substantial amount of lipid uptake, drives intracellular lipid receptor activation through LXR/PPARgamma and induces a transcriptional signature mediating TGFβ and lipid efflux responses in the lung, especially in response to silica particulates (71). CD36 expression in foam cells are increased in patients with silicosis and their presence is associated with intracellular load of oxidized LDL(71). This suggests a role for altered lipid metabolism in this macrophage subset. However, in murine models of lung injury/fibrosis, CD36 promotes apoptotic cell clearance, suppression of inflammation and anti-fibrotic processes in the lung(72). Thus, CD36 may have protective or damaging effects that warrant further investigation.

This study has several limitations. First, our sample size for the CyTOF studies is relatively small. However, the CyTOF panel was specifically designed to address whether we can see an expanded cluster in IPF based on the scRNAseq data, which is the largest of its kind for a human lung with 20 controls, 29 IPF and 18 COPD subjects. Furthermore, the CyTOF findings were corroborated by flow cytometric analysis of biobanked patient samples from the same cohort of subjects, which further affirmed the robustness of the results. Second, our findings linking the identification of CD84 and CD36 as novel surface molecules uniquely mapping an expanded IPF cluster was only performed in end-stage IPF or COPD lungs; thus, we cannot say to what extent this relationship is exclusive to end-stage disease, and how translatable these findings are to stable or progressive disease, since macrophages are highly plastic and their phenotypes likely change over the course of a disease(16). Third, although the evidence showing the expanded IPF cluster in scRNAseq/CyTOF/flow cytometry is robust, we did not elucidate the exact function of the CCL18 producing CD84++, CD36++ macrophage population. Studies have demonstrated that CCL18 is one of the top biomarkers in IPF, predictive of disease progression, and is a key product of pro-fibrotic M2-like macrophages(37,41,73). Although the exact pro-fibrotic role of CCL18 is yet to be fully elucidated, there has been some evidence showing that CCL18 may partially elicit its pro-fibrotic effect on lung resident fibroblasts by recruiting T cells(74). In our study, we did not specifically evaluate fibrosis and therefore, we cannot establish causality of the expanded IPF macrophage cluster to disease pathology. However, based on the presented data and literature evidence, we hypothesize that the reversal of the pro-fibrotic signature in IPF macrophages, combined with the *in vitro* validation demonstrating reduced CCL18 levels from macrophages and cultured IPF PCLS, suggest that the CD84/CD36/CCL18 expressing phenotype might be a pathogenic fibrosis-promoting one. Finally, we used cryopreserved lung digests to allow for batched processing of our specimens to limit experimental noise from day-to-day human and technical variation errors especially with complex techniques that require cellular phenotype evaluation. Our protocol was standardized across all collected lung tissues, but cryopreservation may still have altered certain measurements and results relative to freshly processed cells. Interestingly, a recent report suggests that biobanked, cryopreserved lung cells obtained from human lung explants are viable and serve as valuable resources for large-scale cellular and molecular phenotyping studies(75).

In conclusion, we believe that leveraging scRNAseq/CyTOF/flow cytometry/PCLS is a powerful combinatorial approach for the discovery and validation of a novel IPF-expanded macrophage subset. Our data demonstrate a targetable pro-fibrotic macrophage subpopulation that highly expresses CD84 and CD36, and contribute to enhanced CCL18 levels, suggestive of a pro-fibrotic phenotype. Our study clearly demonstrates the feasibility and utility of high-dimensional single-cell analysis approaches of lung tissues from end-stage IPF/COPD and reveals important insights and strategies to modulate macrophages that may play a role in the development of fibrotic lung disease.

## METHODS

### Human Data availability

De-identified sequencing data for all subjects was download from the gene expression omnibus (GEO) under accession number GSE136831. Data available at www.IPFCellAtlas.com, with different visualization tools, as described (76).

### Lung tissue procurement and single cell suspension and single cell library preparation and sequencing

As previously described(18), a standard protocol for tissue procurement and lung cell isolation was followed(75). Briefly, tissues were obtained from lung explants of subjects diagnosed with IPF and COPD undergoing lung transplantation. Control samples were obtained from rejected lung donors. The study protocols were approved by Mass General Brigham Institutional Board Review (IRB Protocol 2011P002419). Single cell suspensions were obtained after an extensive mechanical and enzymatic tissue dissociation, with multiple filtration steps. Suspensions were cryopreserved in 10% DMSO based media and thawed in batches for furthers steps. Single Cell Library Preparation and sequencing, pre-processing done as specified previously(18).

### Cell Clustering and myeloid cell subset isolation

Highly variable genes using vst method were obtained and later scaled; these values were used as input for PCA. kNN network was built with the first 30 components, with the later generation of cell embeddings using the UMAP algorithm. Cells from Myeloid compartment were selected based on the positive expression of PTPRC gene and negative expression of lymphocyte markers. Harmony was implemented to perform batch correction over subject identity variable (80). Downstream analyses were specifically performed on the myeloid subset. FindAllMarkers function from Seurat package was used to identify cell markers overrepresented in each cluster via the ROC method. Genes were ranked on a descending order based on AUC values.

### PHATE embeddings and pseudo time analysis

PHATE (22) implementation is a dimensionality reduction method that uses diffusion geometry. This method enables better discrimination of the underlying manifold of the data while preserving the local and global distances and the branching progression structure that characterize biological processes. The resulting two-dimensional (2D) visual representation better resembled the ground truth as compared to standard algorithms such as t-distributed Stochastic Neighbor Embedding (t-SNE).

PHATE was implemented on the subset of monocytes and macrophages. 20 kNN, an alpha decay of 5, automated t, and 20 PCAs were the parameters used for PHATE function. Trajectories were then identified using the Slingshot (23)implementation on the PHATE embeddings with default settings. Starting point was the Monocyte cluster. Pseudotime distances were calculated, which identified terminal phenotypes in each of the trajectories.

### Regulon Activity Identification

pySCENIC package was used(24) to score the activity of each regulon using the following databases: cisTarget databases (hg38__refseq r80_500bp_up_and_100bp_down_tss.mc9nr.feather, hg38 refseq-r80__10kb_up_and_down_tss.mc9nr.feather), and the transcription factor motif annotation database (motifs-v9-nr.hgnc-m0.001-o0.0.tbl) and the list of human transcription factors (hs_hgnc_tfs.txt) was downloaded from **github.com/aertslab/pySCENIC/tree/master/resources**.

### Macrophage Polarization scoring

We used Macspectrum (45) algorithm to infer the maturation and polarization of our cells of interest. With collaboration of the publisher team, algorithm for cell-scoring was obtained as an R script and applied to our dataset of myeloid cells. Vectors with the MPI and AMDI were obtained and added to the metadata table of the main Seurat object. Macspectrum performs a linear regression of each cell transcriptional signature and fits it into a pre-specified scRNAseq gene signatures of M1 and M2 macrophages obtained in vitro condition, to then derive a macrophage polarization index (MPI) and an activation-induced macrophage differentiation index (AMDI) per each single cell, which facilitates the classification of cells based on their inflammatory and terminal maturation state

### Implementation of Triwise, radar plots and GO enrichment

Triwise R package (34) was used to perform a comparative differential expression analysis between three final differentiation phenotypes obtained after pseudotime implementation. The three cell types were aggregated by subject identity using the Seurat package(79) function ‘Average Expression’. Then, differential expression using MAST (81) method was implemented to obtain a list of DE genes. Barycentric coordinates were later calculated (34) to create a radar plot as shown in Figure 1D. In this plot, the points are genes, the direction of the point indicates the condition in which the gene is upregulated, and the distance from the origin represents the fold-change. Points lying on a same hexagon grid have the same fold change. Genes that are DE between the three conditions in the same order of magnitude lie at the origin of the plots.

We then calculated the gene ontology enrichment using the GO database and then plot into radar plots, highlight those genes with greater DE amongst each term (Supplemental figure 2). To reduce the redundancy of the gene ontologies, “model-based gene set analysis” method (82) was used, which selected those gene sets that provided optimal explanation for the differentially expressed genes in the dataset.

### Combinatorial prediction of marker panels for IPFeM

*COMETSC* (44) package was used to identified candidate markers for our population of interest. It implements a non-parametric statistical framework to determine gene enrichement in specific population against the rest and true positive and negative rate of each candidate marker. rue Positive rate is found by dividing the number of expressing cells in the cluster by the total cell count of the cluster; true negative is found by dividing the number of non-expressing cells outside the cluster by the total cell count outside the cluster.

### Macrophage Tissue Localization score

We used the bulk-RNAseq dataset from Misharin et al(46), to identify potential origins and tissue locations of our cells of interest. Similarity score per cell was calculated as correlation to differentially expressed genes and log fold changes in the sorted populations. Cells were assigned to either AM or IM category if the score difference for either category is higher than 0.05, according to described prior method(47).

### External Datasets

Correlation plots were created using ggplot2 and complexheatmap(83) R packages, to compare the expression of the macrophage subpopulations identified in our study against the external datasets: Human (48) (GEO accession: GSE135893) and Mouse (47), download (GEO accession: GSE141259)

Ortholog genes with a 1:1 match between the Human and Mouse genome were obtained using getLDS function from BiomaRt package (84) Gene matrix was subset with the ortholog genes, to allow further comparisons.

### Ligand – Target - Receptor Interaction Map

Nichenet R Package was implemented to generate the ligand-receptor analysis. IPFeM were selected as sender cellÍs. We selected Myofibroblast, Aberrant Basaloid Epithelial cells and Vascular Endothelial cells as receivers, to assess the interaction of these macrophages on cells from the fibrotic milieu. Chord plots were then created and labeled according to the main signaling pathways that the L-T pair was part of.

### CyTOF staining and barcoding protocol

Human lung tissue samples were obtained from patients diagnosed with terminal lung disease (IPF and COPD) undergoing lung transplants at the Brigham and Women’s Hospital. Control lungs had no evidence of chronic lung disease and were used as donor controls. Tissues were initially digested into single cell suspensions, as previously described (18,75). Prior to staining, tissue digests were thawed and placed in PBS containing 0.1mg/ml DNAse I solution (Stem Cell technologies, Cat#07900) to digest DNA released from dead cells. Single cell tissue digests were then filtered, counted and processed for subsequent staining and analysis. For CyTOF, 1-3 million cells per subject were prepared for the staining. Initially, viability staining was performed using Cell-ID Cisplatin (5μM in RPMI 1640 with no fetal bovine serum and Penicillin/streptomycin) (Fluidigm, cat#201064) for 2 minutes at room temperature (RT). Next, mild fixation was performed by adding 0.2% paraformaldehyde (PFA) for 5 minutes at RT. Cells were then Fc blocked in Maxpar cell staining buffer (CSB) (Fluidigm, cat#201068) for 10 minutes at RT and then surface antibody staining was performed by adding the metal-coupled Ab cocktail to the cells in CSB (30 minutes at RT). Prior to barcoding each sample, additional fixation was performed using 1X Fix I buffer (Fluidigm) for 10 minutes at RT and cells were then resuspended in 1X barcode perm buffer. Following the barcoding of samples for 30 minutes at RT (Cell-ID 20-Plex PD Barcoding kit, Fluidigm Cat# PN PRD023 A3), all individual samples were combined into one solution tube containing CSB. The sample was further fixed by adding 1.6% PFA solution for 10 minutes at RT. Cells were then stored in 1ml of CSB at 4°C overnight. The following day, MaxPar Cell-ID Intercalator-Ir solution (Fluidigm, cat#201192B) was prepared in Maxpar Fix and Perm buffer (500μM) (Fluidigm, cat#201067) and added to the sample for 20-minutes at RT. Following the incubation period, samples were washed and cell acquisition solution (CAS) was added in addition to EQ four element calibration beads (Fluidigm, cat#201078) for normalization. Cells were counted and normalized to a final concentration of 0.75 x 10^6^ cells/ml before CyTOF analysis. Samples were run using the Helios CyTOF system.

### CyTOF analysis

CyTOF analysis was performed as previously described (51). Briefly, mass cytometry data was analyzed using Cytobank online software (https://cytobank.org) to perform t-Distribution Stochastic Neighbor Embedding (tSNE) analysis(85). Optimal clustering density was determined by X-shift algorithm using serial iterations of K-nearest neighbor estimation. A range of optimal cluster numbers was used to run SPADE analyses to group phenotypically similar cell populations. Differentially abundant clusters were identified using one-way analysis of variance with Bonferroni’s multiple comparisons test.

### Flow Cytometry staining and analysis

Independent cohorts (N=26, 6 healthy controls, 11 IPF and 9 COPD) were included for additional validation. Flow cytometry panel design and gating strategies were adapted from published work phenotyping cells derived from human lung tissue and bronchoalveolar lavage fluid(86),(54). Cells were initially stained with a Zombie live/dead viability dye (Biolegend, Cat#423101) in phosphate buffered saline for 30 minutes at RT. Next, samples were washed with FACS buffer (0.3% BSA in PBS) and stained with Human TruStain FcX (BioLegend, Cat# 422301) for 15 minutes. Samples were subsequently stained with the antibody cocktail mix in FACS buffer containing, anti-human CD45 (APC Fire 750, Biolegend, Cat#368518), anti-human HLA-DR (PerCp Cy5.5, BD Pharmingen, Cat#560652), anti-human CD15 (PE Cy7, Biolegend, CaT#323030), anti-human CD11b (AF700, Biolegend, Cat#101222), anti-human CD169 (BV605, Biolegend, Cat#346010), anti-human CD206 (BV421, BD Pharmingen, Cat#564062), anti-human CD36 (APC, Biolegend, Cat#336208) and anti-human CD84 (PE, Biolegend, Cat#326008) for 30 minutes in 4 degree Celsius. Finally, samples were washed twice, resuspended in FACS buffer, and analyzed using a BD LSRFortessa flow cytometer. Data were analyzed using FlowJo version 10.2. Cells were sequentially gated on single cells ((FSC-A vs. FSC-H) and (FSC-W vs. FSC-A)), viable cells (Zombie negative) and immune cells (CD45+ subset). Neutrophils were excluded by gating on the CD15-compartment and the monocyte/macrophage gate was selected (CD11+HLADR+ subset). By excluding monocytes (CD206-CD169-subset), both alveolar and interstitial macrophages were selected based on CD206 and CD169 expression. The assessment of CD84 and CD36 expression was performed on the macrophage gate using fluorescent-minus-one controls

### Generation and treatment of THP-1-derived macrophages

The THP-1 human monocytic cell line was purchased from the American Type Culture Collection (ATCC#TIB-202). These suspended cells were grown in RPMI-1640 medium supplemented with 2 mM L-glutamine, 1% Penicillin Streptomycin and 10% FBS. THP-1 monocytes were differentiated into macrophages using Phorbol Myristate Acetate (PMA) (ATCC-Cat#202152) at 10 ng/ml for 48 hours. Macrophages were then skewed towards the M2 phenotype using recombinant human IL-4 (20 ng/ml) (Peprotech, Cat#AF-200-04), IL-13 (20 ng/ml) (Peprotech, Cat#AF-200-13), recombinant human IL-6 (10 ng/mL) (Peprotech, Cat#AF-200-06) and where applicable, drugs were added. Cucurbitacin I (Cucur; Cayman Chemicals, Cat#14747), Cinobufagin (APExBIO, Cat#N1154) and Cryptotanshinone (CRT; Millipore Sigma, Cat#35825-57-1). All drugs were prepared in the solvent dimethyl sulfoxide (DMSO) and all conditions were adjusted to a final concentration of 0.1% DMSO. Exposure to the above cocktails and drugs lasted for 72 hours before collection of supernatant and RNA isolation. Cellular viability/toxicity of the THP-1 derived macrophage system was established with the MTS cellular proliferation-viability assay (Promega, Cat#G3582).

### ELISA

Human CCL18 and TNFα protein was assessed in the cell culture supernatant using the commercially available CCL18 and TNFα ELISA, according to the manufacturer’s protocol (R&D systems, Cat# DCL180B and Cat#DY210-05).

### IPF precision cut lung slices (PCLS)

Human IPF lungs were obtained from patients with terminal fibrotic lung disease undergoing lung transplantation at Brigham and Women’s Hospital. All experimental procedures were performed under sterile conditions and the protocol was adapted from a previously described paper(87). Upon proper cannulation of the donor IPF lung, pre-warmed low melting dose of agarose (Thermofisher Scientific, Cat#16520050) was injected into the lung through the mainstem bronchus until the lung was fully inflated. The inflated lung was placed on ice for 30 minutes to solidify the agarose. A disposable biopsy punch was then used to create tissue cores 100mm in diameter. To facilitate the embedding and sectioning process, the tissue core was glued to the specimen tube which was then filled with warm agarose. Lung sections (100mm diameter, 350μm thick) were prepared with VF-300-0Z Vibratome (Precisionary instruments, Natick, MA).

Tissue sections were then cultured in media (RPMI1640 plus 10%FBS and 1%P/S) containing cryptotanshinone and cucurbitacin (100nM) for 48 hours. After 48 hours, the slices were either processed for digestion and flow cytometry staining/analysis or saved in TriZol RNA extraction using the TriZol method. The PCLS supernatant was saved for ELISA.

### RNA extraction and assessment of RNA quality

Total RNA from THP1-derived macrophages was isolated using Nucleospin RNA plus (MACHEREY NAGEL) according to the manufacturer’s protocol. Total RNA from IPF PCLS was isolated using the traditional TRIzol method according to the manufacturer’s protocol (Thermofisher Scientific, Cat#15596018). Trizol-isolated RNA was then subjected to DNase treatment with Deoxyribonuclease I (Life technologies, Cat#18068-015).

### Real-time polymerase chain reaction (RT-PCR)

RNA isolated from THP1-derived macrophages and IPF PCLS were reverse-transcribed using Superscript IV Reverse transcriptase (Thermofisher, Cat#18090050) to obtain cDNA for gene expression analysis. A Bio-Rad Real-Time PCR Machine with iTaq Universal SYBR Green Supermix (BioRad Catalog#1725122) were employed. The PCR protocol used was a 20-second polymerase activation and DNA denaturation at 95°C, followed by a 2-second denaturation at 95° C, a 15-second annealing/extension and plate read at 60°C, 40 cycles. SYBR green primers, including *Ccl18*, forward, CTCCTTGTCCTCGTCTGC, and reverse, CTATGAACTTTTGTGGAATCTGCC and for *18S*, forward, ACATCGCTCAGACACCATG, and reverse, TGTAGTTGAGGTCAATGAAGGG were produced by Integrated DNA Technologies IDT. *18s* was used as reference gene to assess *Ccl18* mRNA gene expression. Candidate genes were analyzed using semi-quantitative gene expression analysis (ΔΔCT method) and expressed as fold change relative to the gene expression of the control untreated condition.

