## Supplementary Material for "Single Cell RNA-seq and Mass Cytometry Reveals a Novel and a Targetable Population of Macrophages in Idiopathic Pulmonary Fibrosis"

**Supplementary Figure 1:**

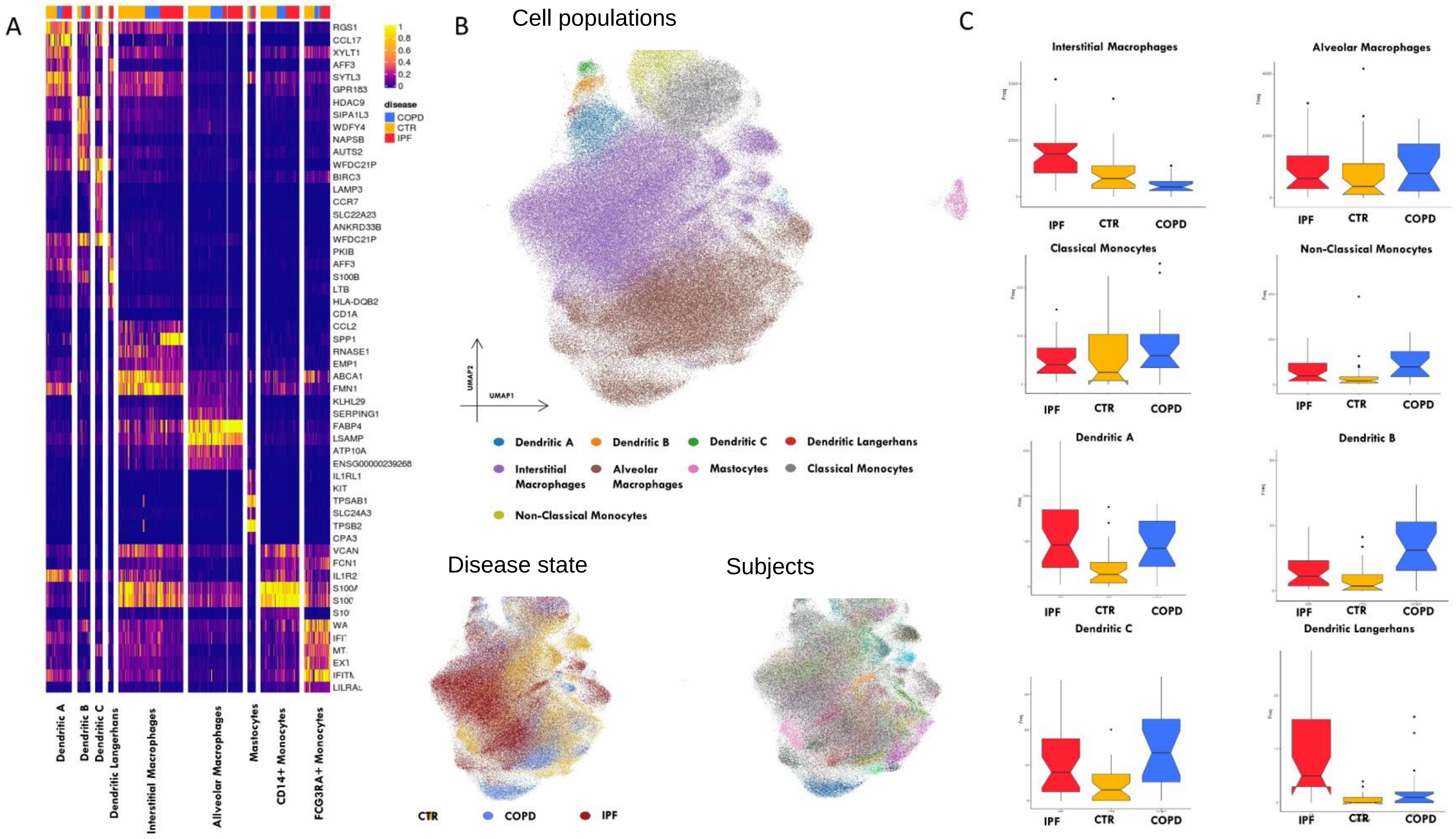

**Supplementary Figure 1:** Myeloid compartment and cell subpopulations. A. Heatmap with DE markers for each subpopulation. Cell populations are labeled with disease state annotation in the horizontal bar (COPD [blue bars], Control [CTR; orange bars], and IPF [red bars]). B. UMAP plots of the entire myeloid compartment, colored by cell population, condition, and subject identities in each plot. C. Violin plots with compositional analysis of each subpopulation.

**Supplementary Figure 2:**

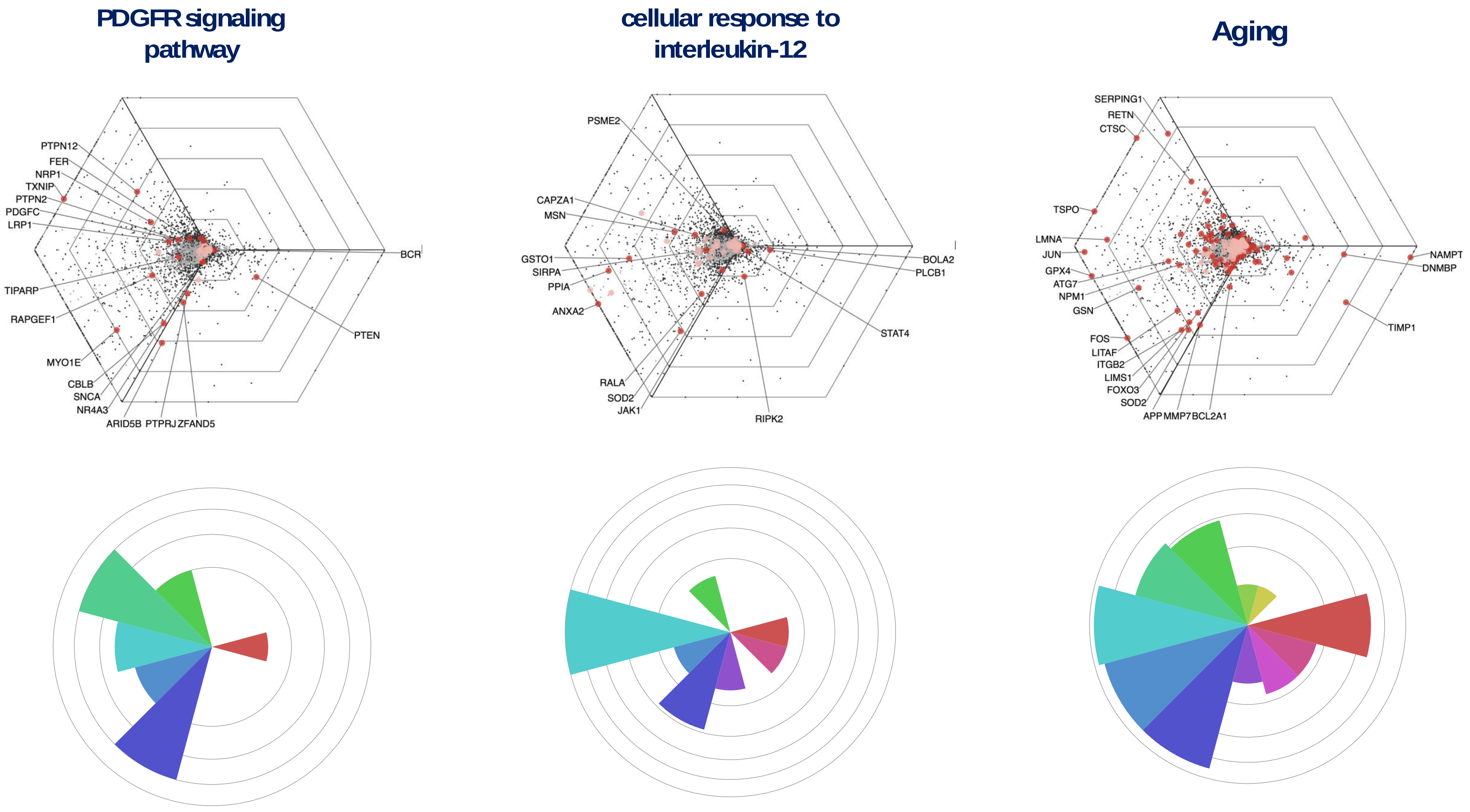

**Supplementary Figure 2: Triwise radar and corresponding polar plots of PDGF, IL-12, and aging signaling pathways.**

**Supplementary Figure 3:**

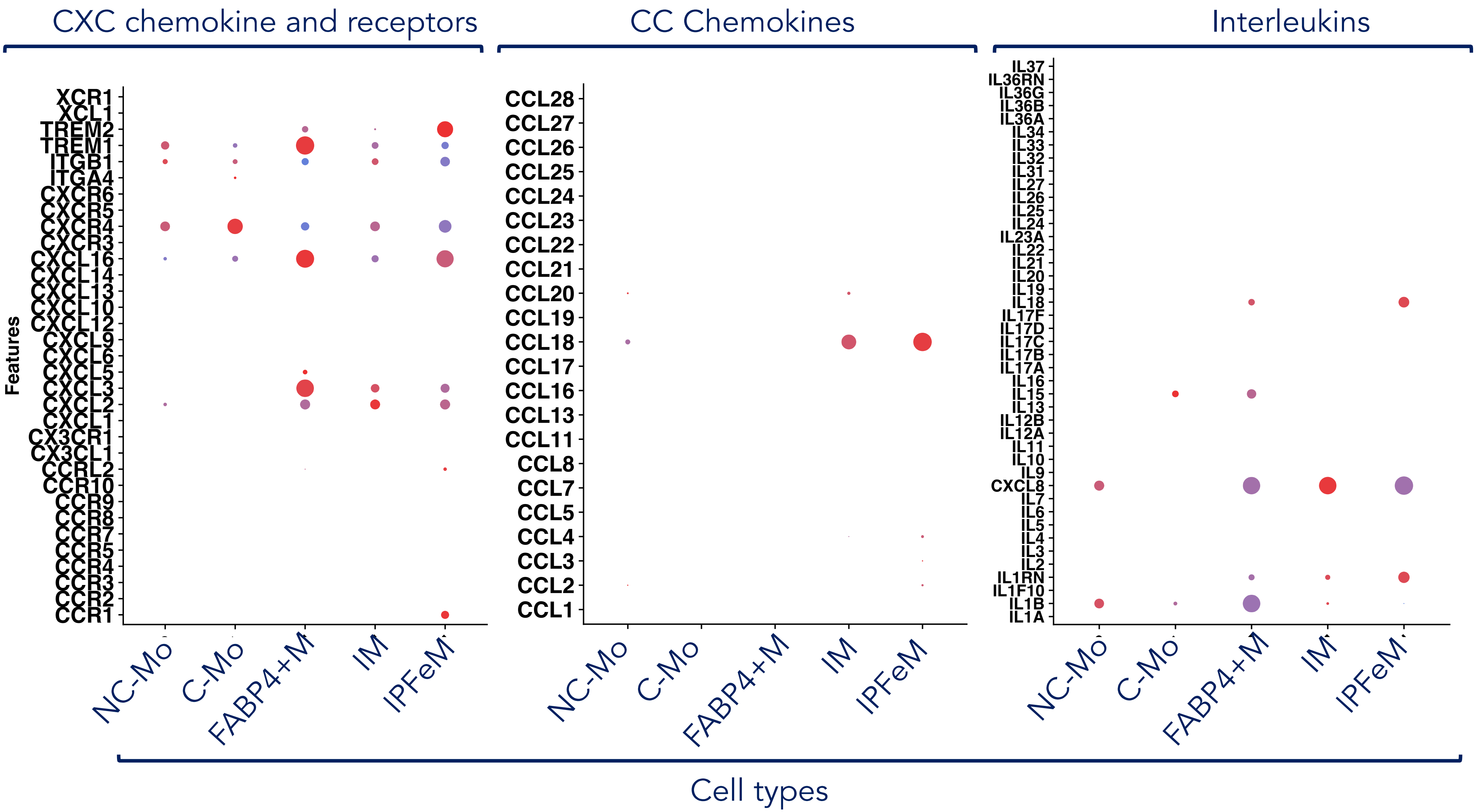

**Supplementary Figure 3: Dot Plot visualization of Cytokine-related genes**, including: CXC chemokines, CC Chemokines, Interleukins and chemokine receptors. Monocyte/Macrophage cell types depicted, NC-Mo: Non-Classical Monocyte, C-Mo: Classical Monocyte, FABP4+M: FABP4 expressing Macrophage, IM: Interstitial Macrophage, IPFeM: IPF - expanded Macrophage. Interleukins most expressed by the IPFeM are IL1RN, CXCL8 and IL18; FABP4+M had overexpression of IL1B, CXCL8, IL15. CC chemokine mostly expressed by IPFeM was CCL18, which had the highest expression amongst myeloid cells. CXC chemokines and chemokine receptors expressed by the IPFeM were CCR1, CXCL2, CXCL3, CXCL16, and TREM2. FABP4+M shown overexpression of TREM1, CXCL16, CXCL3.

**Supplementary Figure 4:**

### Cell Surface Singletons

### True negative vs True Positive curve, for IPFeM cluster

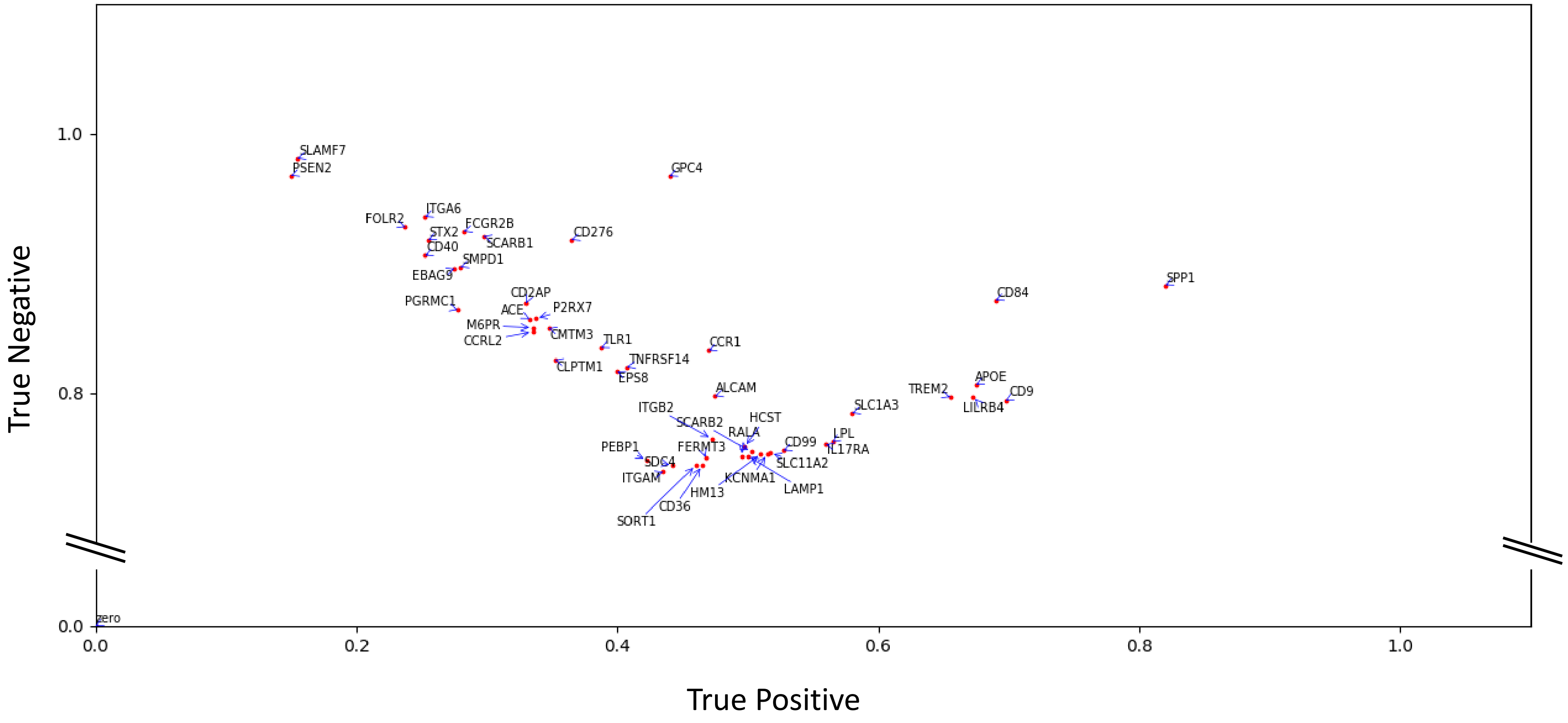

**Supplementary Figure 4:** True positive/true negative discriminatory ratio for IPFeMΦ cells of different cell-surface protein coding genes, calculated by COMETSC package. Both x and y axis are scaled percentages, 1.0 corresponds to 100 percent. SPP1, followed by CD9, CD84, APOE, LILRB4 and TREM2 are the genes with the highest true positive values, whereas SLAMF7, PSEN2 and GPC4 are the genes with the highest true negative values.

**Supplementary Figure 5:**

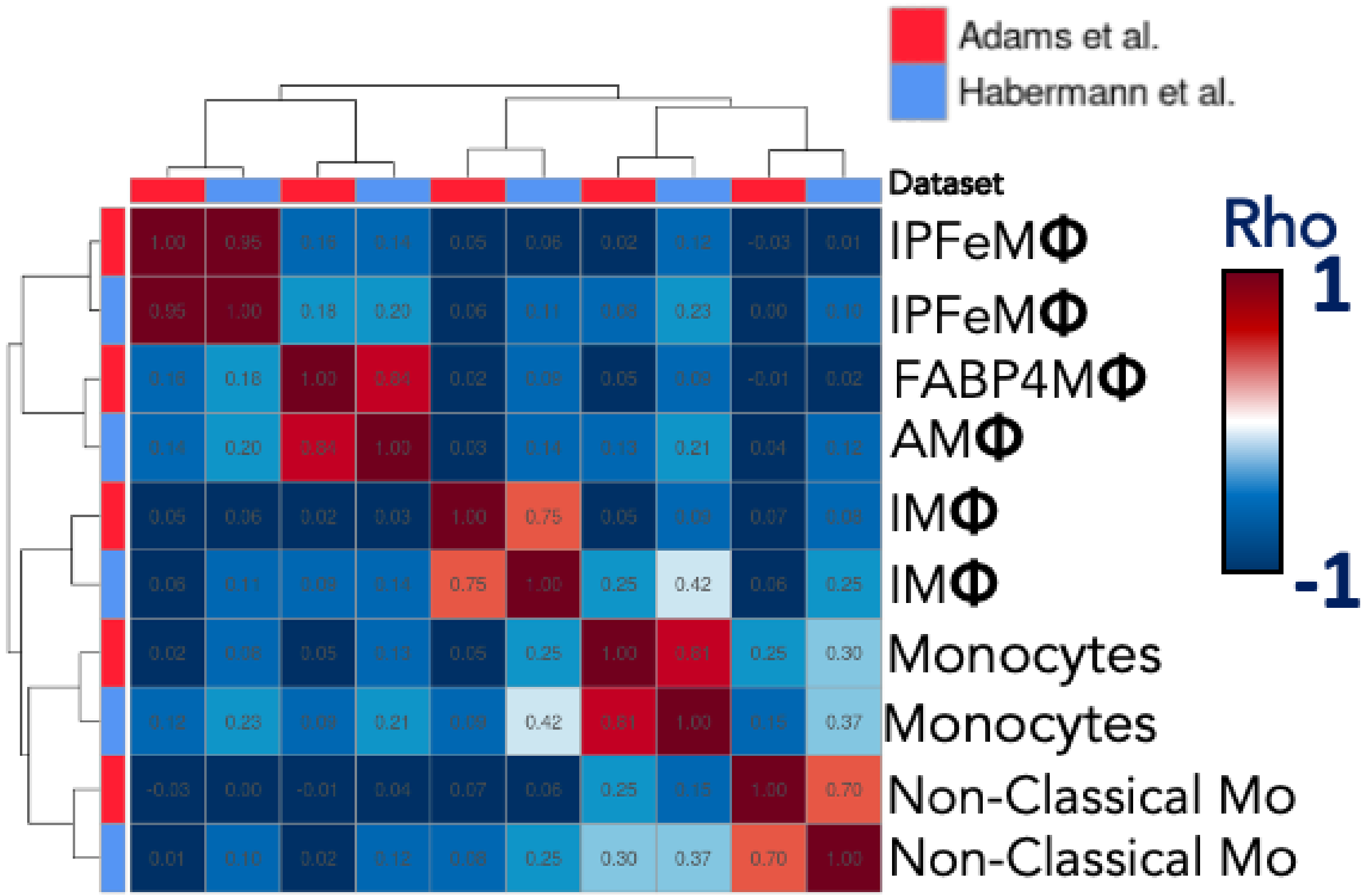

**Supplementary Figure 5: Correlation plot using myeloid cells from Habermann dataset evidenced a similar population (rho = 0.95) of cells expanded in fibrotic lungs.**

**Supplementary Figure 6:**

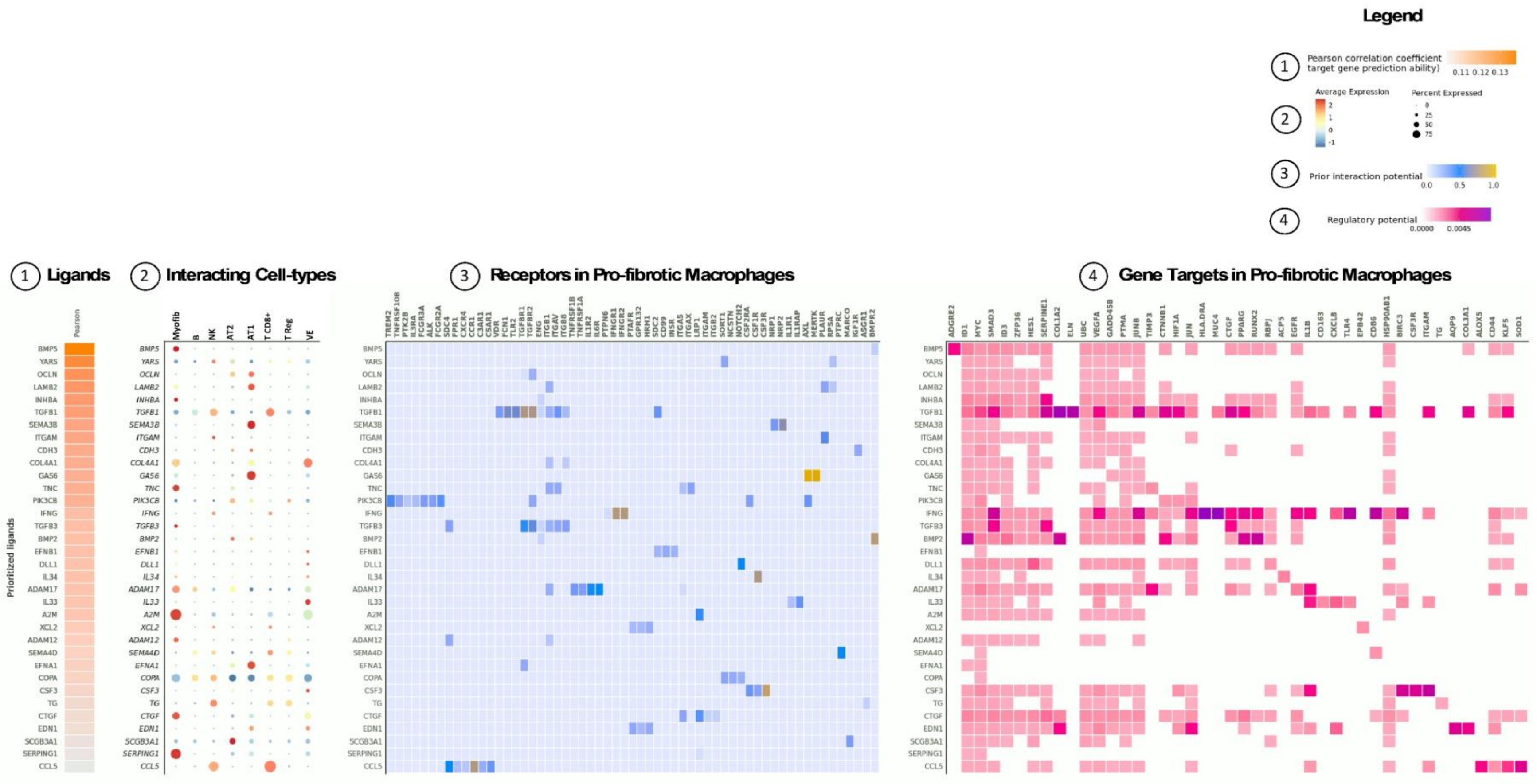

**Supplementary Figure 6: Ligand Receptor Interaction Map of IPF lung, pro-fibrotic macrophages interactions.** NicheNet analysis of Pro-fibrotic Macrophages depicting 1) top ligands prioritized by Pearson correlation analysis. 2 is showing the cell type specificity of the prior ligands on different cell subtypes that are interacting with the macrophage. 3, Receptors in Profibrotic Macrophages that interact with the prioritized ligands. 4, Downstream targets in pro-fibrotic macrophages, regulated by the prioritized ligands on different cell subtypes.

**Supplementary figure 7**

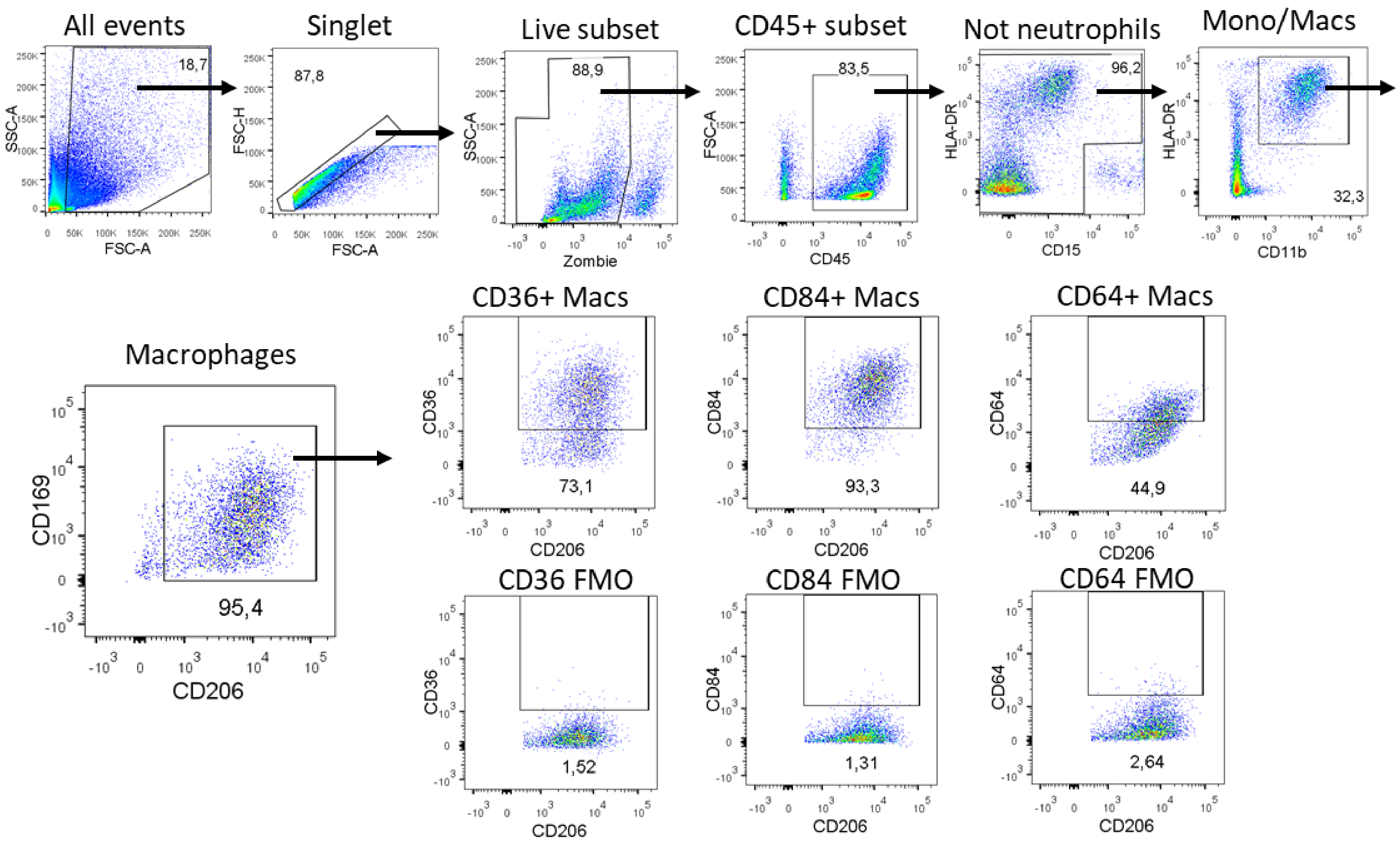

**Supplementary figure 7.** End-stage IPF lung was inflated with low melting dose agarose and precision cut lung slices were cultured in media. After 48 hours, tissues slices were subjected to flow cytometric analysis. A gating strategy showing cellular viability (Zombie (Live) – subset) and detailed sequence of flow plots leading to the expression of CD36, CD84 and CD64 within the combined macrophage compartment (alveolar and interstitial macrophages)

**Table 1**

| Target | Label | Target | Label |
|---|---|---|---|
| CD41 | 089Y | FOLR2 | 159Tb |
| CD45 | 115In | CD59-p282 | 160Gd |
| Syndecan 2 | 141Pr | CD66a | 161Dy |
| CX3CR1 | 142Nd | CD8a | 162Dy |
| CD123 | 143Nd | CD33 | 163Dy |
| CD69 | 144Nd | CD15 | 164Dy |
| CD217 | 145Nd | CD200R1 | 165Ho |
| CD64 | 146Nd | CD24 | 166Er |
| CD11c | 147Sm | CD11b | 167Er |
| CD116 | 148Nd | CD206 | 168Er |
| GARP | 149Sm | IL-4R | 169Tm |
| CD369 | 150Nd | CD3 | 170Er |
| CD14 | 151Eu | CD20 | 171Yb |
| CD36 | 152Sm | CD38 | 172Yb |
| CD192 | 153Eu | HLA-DR | 173Yb |
| CD163 | 154Sm | CD84 | 175Lu |
| IGFR2 | 155Gd | CD4 | 176Yb |
| CD204 | 156Gd | CD16 | 209Bi |
| CD169 | 158Gd | | |

**Table I. CyTOF heavy metal conjugated antibodies**
